# A proposed function for the red body of *Nannochloropsis* in the formation of the recalcitrant cell wall polymer, algaenan

**DOI:** 10.1101/2023.09.17.558137

**Authors:** Christopher W. Gee, Johan Andersen-Ranberg, Rachel Z. Rosen, Danielle Jorgens, Patricia Grob, Hoi-Ying N. Holman, Krishna K. Niyogi

## Abstract

Stramenopile algae contribute significantly to global primary productivity, and one class, Eustigmatophyceae, is increasingly studied for applications in high-value lipid production. Yet much about their basic biology remains unknown, including the nature of an enigmatic, pigmented globule found in vegetative cells. Here, we present an in-depth examination of this “red body”, focusing on *Nannochloropsis oceanica*. During the cell cycle, the red body formed adjacent to the plastid, but unexpectedly it was secreted and released with the autosporangial wall following cell division. Shed red bodies contained antioxidant ketocarotenoids, and overexpression of a beta-carotene ketolase resulted in enlarged red bodies. Infrared spectroscopy indicated long-chain, aliphatic lipids in shed red bodies and cell walls, and LC-HRMS detected a C32 alkyl diol, a potential precursor of algaenan, a recalcitrant cell wall polymer. We propose that the red body transports algaenan precursors from plastid to apoplast to be incorporated into daughter cell walls.

## Introduction

### Introduction to eustigmatophyte algae

The class Eustigmatophyceae is a clade of stramenopile microalgae that has been the subject of a growing body of work in the past decade, due in part to their rapid growth and ability to partition large fractions of their biomass into valuable lipids such as the omega-3 fatty acid, eicosapentaenoic acid (EPA) (Chaturvedi and Fujita, 2006; Dolch et al., 2017; Sukenik, 1990). Algae from this class were reputed to be uncommon, but recent phylogenetic reassessments have yielded a much expanded tree (Amaral et al., 2020; Fawley et al., 2014; Fawley and Fawley, 2017), and eustigmatophyte algae have been isolated from freshwater, marine, and terrestrial environments around the world (Eliáš et al., 2017; Gao et al., 2019, 2016; Jo and Hur, 2015).

In recent years, the marine nanophytoplankton *Nannochloropsis* and its sister genus *Microchloropsis* (Fawley et al., 2015), have been established as model systems. Cells are typically solitary, non-motile coccoids 2-4 μm in diameter (Fawley and Fawley, 2007), and they reproduce by asexual fission during the night period (Poliner et al., 2015; Sukenik and Carmeli, 1990). Yields for *Nannochloropsis* lipid content have been reported up to ∼50% of the biomass total (Ma et al., 2014), and much research has been directed towards developing elite lipid-producing strains (Jinkerson et al., 2013; Ryu et al., 2020; Wang et al., 2014) and optimizing culture conditions (Simionato et al., 2013; Zienkiewicz et al., 2020). Molecular genetic studies of these genera have been facilitated by the publication of reference genomes (Corteggiani Carpinelli et al., 2014; Radakovits et al., 2012; Vieler et al., 2012) and the development of gene editing tools (Kilian et al., 2011; Poliner et al., 2019, 2018; Wang et al., 2016). Additionally, basic research has highlighted the potential of these algae for providing insights into algal CO_2_-concentrating mechanisms (Gee and Niyogi, 2017; Huertas et al., 2000), photosystem I supercomplex structure (Bína et al., 2017), light-harvesting pigments (i.e., violaxanthin and vaucheriaxanthin) (Keşan et al., 2016), non-photochemical quenching (Carbonera et al., 2014; Park et al., 2019), cell division (Murakami and Hashimoto, 2009), and endosymbiosis with a novel clade of *Phycorickettsia* (Ševčíková et al., 2019; Yurchenko et al., 2018, 2016).

### The red body of Eustigmatophyceae

A diagnostic characteristic of eustigmatophyte vegetative cells is the presence of a red-orange globule that resides outside of the chloroplast (Eliáš et al., 2017). This feature has been observed in several eustigmatophytes (Amaral et al., 2020; Gao et al., 2019; Nakayama et al., 2015; Přibyl et al., 2012), including *Nannochloropsis* specifically (Antia and Cheng, 1982; Fawley et al., 2014; Fawley and Fawley, 2007; Suda et al., 2002). The structure has been variously referred to as a “lipid body”, “reddish globule”, “pigmented spherule”, “eyespot”, “red body”, etc. The original description of the class by Hibberd & Leedale established Eustigmatophyceae by extricating species from Xanthophyceae based on zoospore morphology that included an unusual extraplastidial anterior eyespot, which we presume to be the eponymous “stigma” (Hibberd and Leedale, 1970). Hibberd later authored an update on the classifications of Eustigmatophyceae and included a description of a “reddish globule” in vegetative cells that was different from the “red extraplastidial eyespot” in zoospores (Hibberd, 1981). Given the relatively common use of “red body” in existing work to refer to the red globular feature in vegetative cells, and that this appears to be distinct from the zoospore eyespot, we will refer to it here as the “red body”.

Despite the apparently widespread occurrence of the red body across Eustigmatophyceae, no in-depth examinations have been performed on it beyond a simple description of size and color from microscopy. To our knowledge, no detailed account of its formation has yet been reported, nor a function proposed.

### Summary

Here we present, to our knowledge, the first in-depth examination of the eustigmatophyte red body. We focused on the model species, *Nannochloropsis oceanica*, and characterized the development of the red body over the course of the cell cycle by light and electron microscopy. The red body was secreted during cell division, and shed into the medium along with the autosporangial wall, which facilitated its isolation and enrichment for subsequent mass and infrared spectroscopy and chromatography. These analyses indicated the presence of ketocarotenoids and long-chain aliphatic lipids. We propose that the red body accumulates lipidic precursors of the polymer algaenan, along with ancillary proteins, and upon autospore maturation, delivers these molecules into the apoplast where they polymerize to form the recalcitrant outer layer of the daughter cell walls. Implications for future research of eustigmatophyte algae and understanding the biosynthesis of hydrophobic biopolymers in general are discussed.

## Results

### The red body of eustigmatophyte algae is an autofluorescent, globular, membrane-bound organelle

In our previous studies of *Nannochloropsis oceanica* CCMP1779 (Gee and Niyogi, 2017), we observed unexpected autofluorescent punctae that were distinct from the plastid. Although the physical origin of the emitted light from these punctae remains unknown, we will refer to it as autofluorescence. By transmitted light microscopy, a corresponding reddish globule was sometimes visible. We obtained other eustigmatophyte species and again observed punctate, extra-plastidic fluorescence that corresponded with a red body (Figure 1). The red body autofluorescence was visible through filter sets spanning most of the visible spectrum (Figure 1, supplemental 1). Additionally, for *Chloridella neglecta* (SAG 48.84), confocal scanning laser microscopy revealed that the red body was an aggregation of smaller bodies in this species (Figure 1, supplemental 2). Green autofluorescence (GAF) from the red body was generally robust and spectrally distinct from red chlorophyll fluorescence, so subsequent microscopy used settings similar to those for green fluorescent protein (excitation 488 nm, emission 495-550 nm). The well established eustigmatophyte model organism, *Nannochloropsis oceanica*, was selected for subsequent experiments.

**Figure 1.**
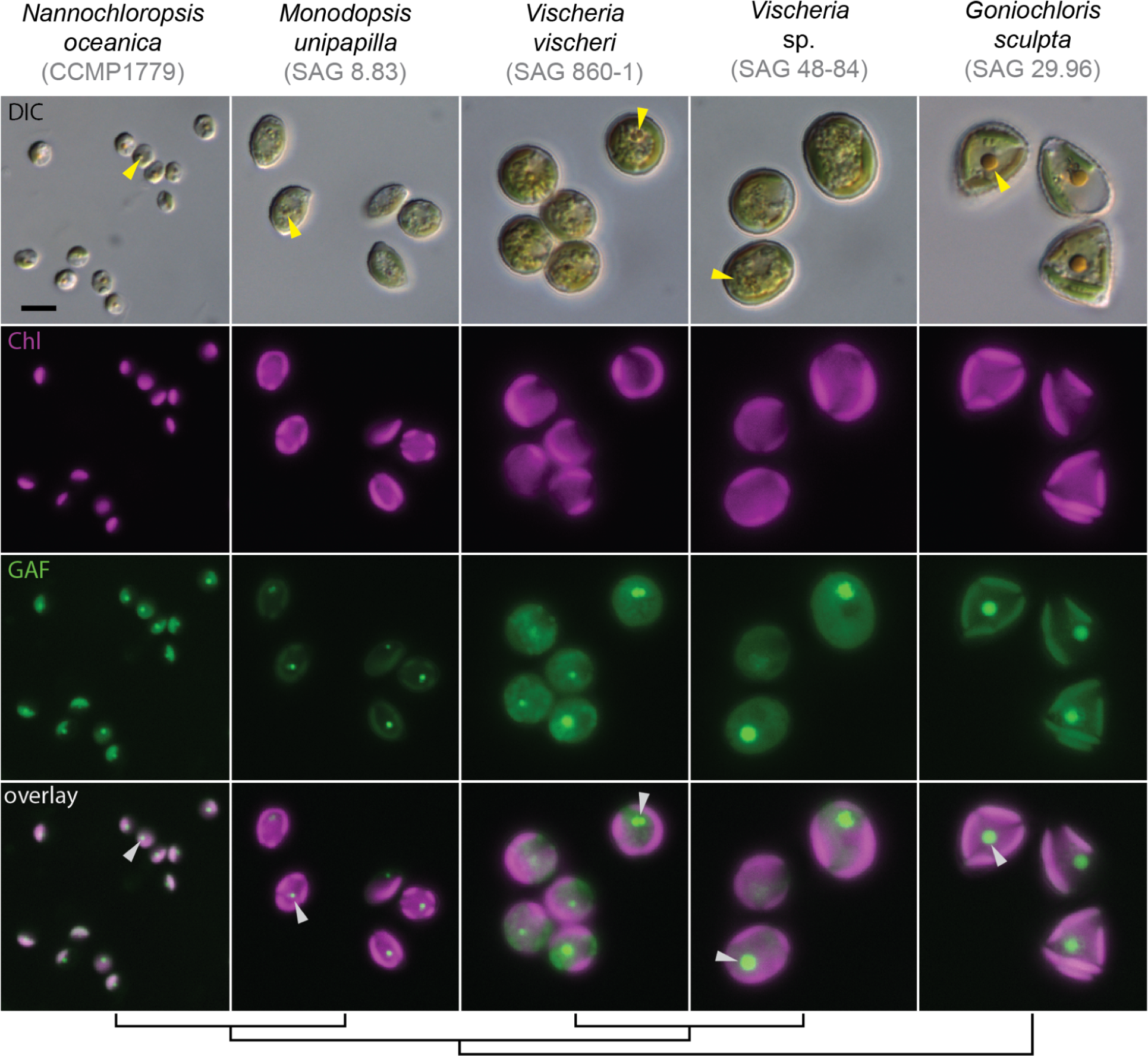
The red body is a globular, autofluorescent subcellular structure found in eustigmatophyte algae. Wide-field fluorescence microscopy. From top row to bottom: differential interference contrast (DIC), chlorophyll autofluorescence (Chl-pseudo colored magenta, ex/em: 660 nm / 700 nm), green autofluorescence (GAF-pseudo colored green, ex/em: 470 nm / 525 nm), overlay of Chl and GAF. An example red body is indicated for each species by an arrowhead (yellow for DIC images, white for overlay). The cladogram at the bottom is based on Amaral et al., 2020. The scale bar in the upper left image equals 5 μm and applies to all images in this figure.

Transmission electron microscopy (TEM) revealed additional details of red body morphology. Chloroplasts served as landmarks and were readily identified by thylakoids grouped in stacks of three, which is typical of secondary plastids derived from red algae (Flori et al., 2017). Based on our observations by fluorescence microscopy, we identified candidate structures in TEM of thin sections that appeared likely to be the red body based on size, shape, and relative position to the chloroplast. These candidate red bodies were circular, ∼300-400 nm in diameter, homogenous and moderately electron dense, and often located adjacent to the chloroplast (Figure 2). At least one membrane was clearly visible encompassing the red body (Figure 2), and in some cases this was enclosed by another membrane encircling the chloroplast (Figure 2A). This enclosing membrane likely corresponds to the outer membrane of the chloroplast-endoplasmic reticulum (CER), a contiguous membrane network formed by the outer nuclear envelope, ER, and the outermost plasmid membrane (Gibbs, 1979; Murakami and Hashimoto, 2009).

**Figure 2.**
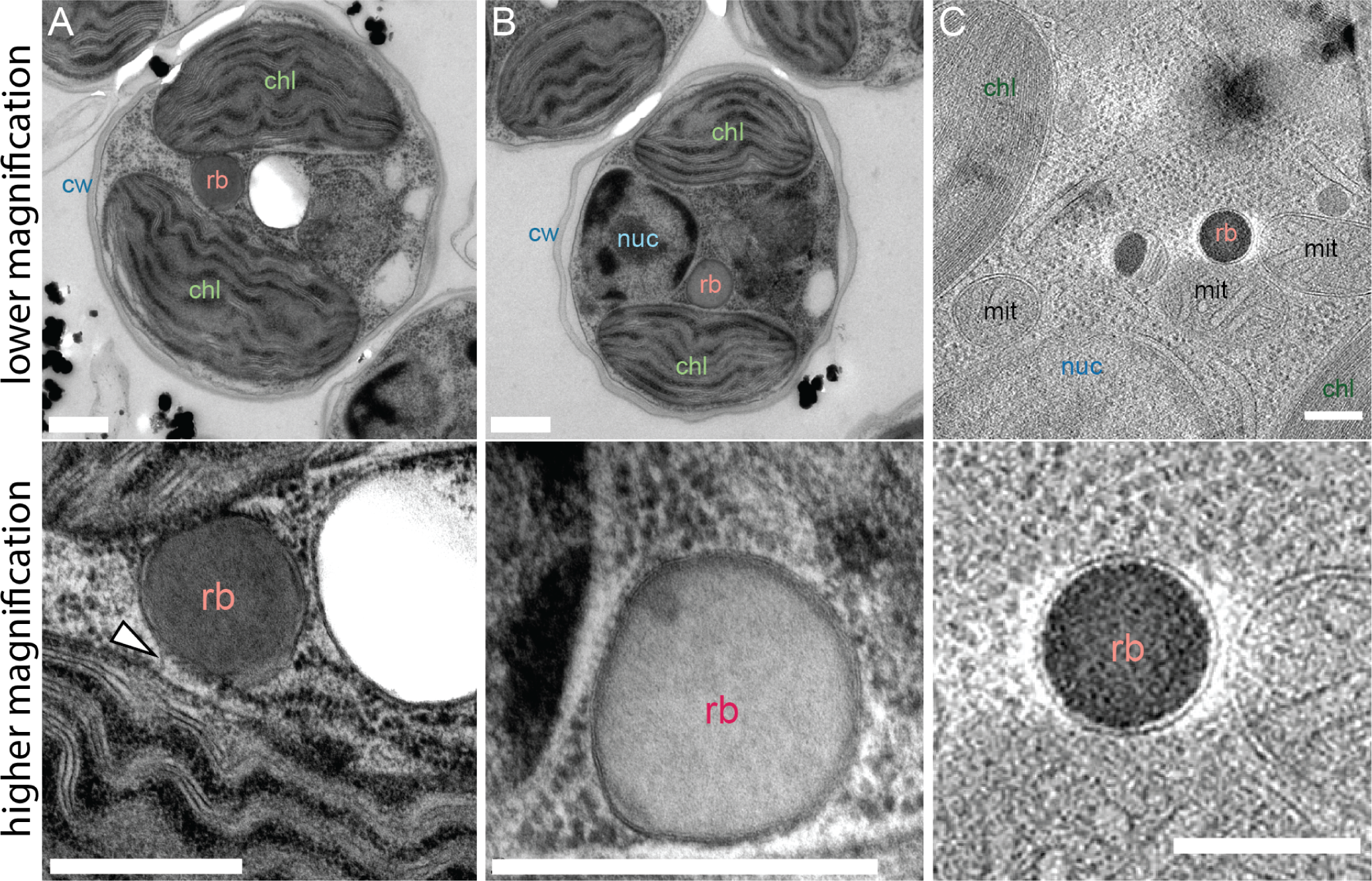
A membrane envelopes the red body. **(A) + (B)** Resin embedded, heavy metal stained transmission electron micrographs of two separate *Nannochloropsis oceanica* cells preserved by high-pressure freezing at subjective dusk. Presumptive assignments: cw = cell wall, chl = chloroplast, rb = red body, nuc = nucleus. The lower image was acquired at higher magnification. A possible membrane connecting the red body with the plastid is indicated with a white arrowhead in (A). **(C)** A selected slice from the cryo-tomogram capturing the red body with apparent enveloping membrane (whole series available as Video 1). mit = mitochondrion. The lower panel is a digitally enlarged portion of the upper image. For all images, the scale bar = 500 nm.

To complement observations made by traditional resin-embedded TEM, cryo-electron tomography was employed to investigate native-state ultrastructure of the red body within *Nannochloropsis*. Cryo-fluorescence microscopy was used to guide cryo-focused ion beam (FIB) milling of thin lamellae for cryo-electron tomography (Figure 2, supplemental 1). The tomogram (Video 1) contained a rich collection of cellular features. These include a spherical red body, clearly enveloped by a membrane (Figure 2C). Additionally, a great many other cellular features were visible, and the preserved detail of these structures highlights the promise of this technique for future investigations of *Nannochloropsis* cell biology. Given the clear compartmentalization and differentiation of its contents from the rest of the cell, we propose that the red body be defined as a membrane-bound organelle.

### Development of the red body is integrated into the cell cycle of *N. oceanica*

In cells grown under continuous light, the size and position of the red body was variable. Growing cells under a diurnal photoperiod (12 hours light, 12 hours dark) not only induced synchronous cell division but also revealed that the red body exhibits regular changes in size and position over the course of the cell cycle. *N. oceanica* cells enlarge during the day period, divide, then separate into daughter autospores during the night period (Poliner et al., 2015) (Figure 3A). To improve resolution of these small features, we used super-resolution structured illumination (SIM) fluorescence microscopy at timepoints throughout the day/night cycle. During the day period, chloroplasts (red fluorescence) grew from flattened ellipsoids to bi-lobed shapes which divided around the light-dark transition to form two daughter plastids (Figure 3B, C). These subsequently divided again to form four plastids, presumably held within four daughter autospores, indicating that *Nannochloropsis* can divide by multiple fission as seen in some other algae (Bišová and Zachleder, 2014). The red body (green fluorescence) began to be clearly visible by ∼4 hours into the light period as small, round punctae next to the plastid. These continued to grow adjacent to the chloroplast, but later moved to the cell periphery during the night (Figure 3B, C). Newly separated autospores contained one plastid but no visible red body, suggesting that it is “shed” upon autospore separation and is subsequently generated anew in each daughter cell.

**Figure 3.**
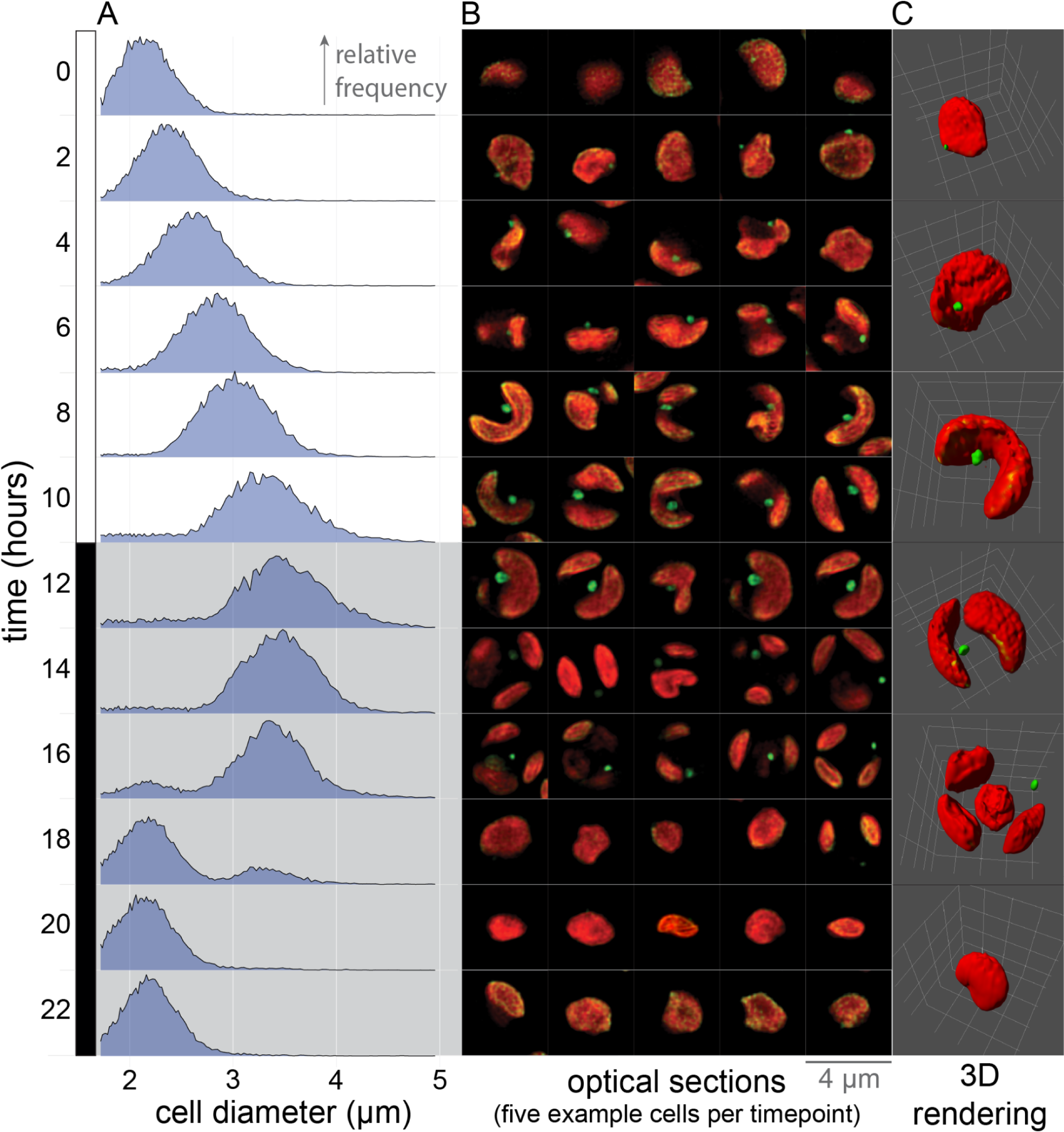
The formation of the red body is integrated into the diurnal cell cycle of *Nannochloropsis*. **(A)** A liquid culture of synchronously dividing cells was sampled at the indicated time points (0 hr = lights on), and cell diameter was quantified by Coulter counter. Resulting distributions are plotted as relative frequency with the same vertical scaling (bin size ∼ 0.01 μm, maximum vertical height ∼3.6%, for each distribution, n > 10k). **(B)** SIM optical z-sections of cells sampled at the same time as in (A). Chlorophyll autofluorescence (excitation 642 nm, emission > 655 nm, pseudo colored red); green autofluorescence (excitation 488 nm, emission 495 to 550 nm, pseudo colored green). Z-position for each cell was chosen to maximize the apparent diameter of the red body. Each individual cell image frame measures 4 x 4 μm. **(C)** 3D reconstructions derived from SIM z-stacks of cells chosen to represent stages of the cell cycle. Each 3D rendering is bounded by a 4 x 4 x 4 μm volume, demarcated on the edges of the volume by a white grid.

### The red body is secreted into the apoplastic space prior to autospore release

To test this idea, we immobilized synchronized cells onto polylysine-coated microscope coverglasses and observed what was left adhered to the glass after cell division and separation. Initial attempts at live-cell imaging over the entire timeframe were unsuccessful, possibly due to cellular sensitivity to light during division, so pairs of coverglasses were made from the same culture and imaged before or after autospore separation (Figure 4A, B). Green autofluorescent punctae were observed adhered to the glass after autospore separation, sometimes trapped within a transparent, pointed, tubular structure (Figure 4B). We propose that these structures are shed autosporangial walls that previously contained the growing autospores during maturation. The released autospores were then bound onto new coverglasses, and no red body punctae were observed (Figure 4C), again indicating that the structures are formed de novo in each daughter cell.

**Figure 4.**
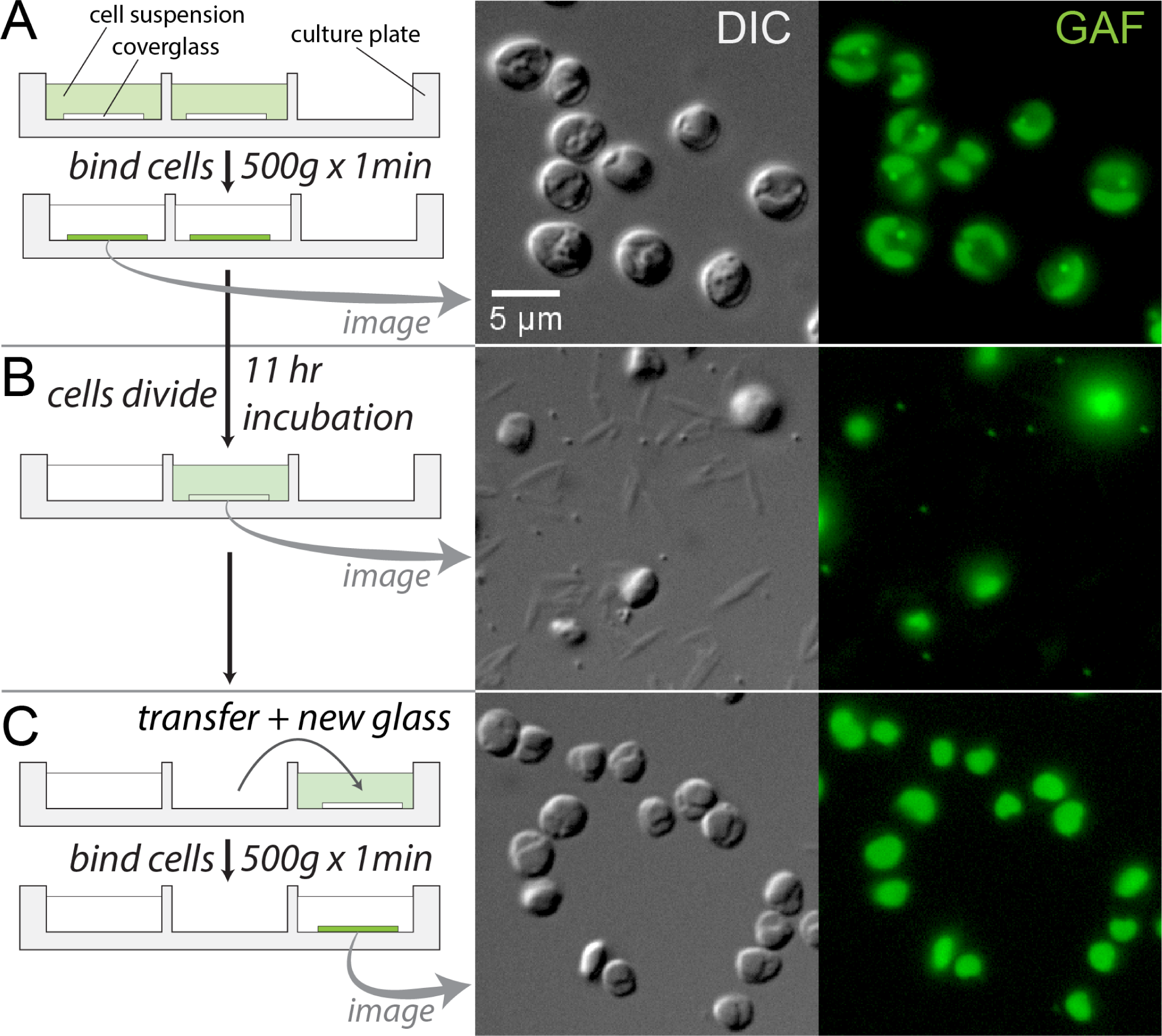
The red body is shed with the autosporangial wall upon autospore release. **(A)** Aliquots of a synchronously dividing liquid culture entrained to a 12 hour light, 12 hour dark photoperiod were bound at subjective dusk to polylysine-coated microscope coverslips by centrifugation in cell culture plate wells. The first coverslip was imaged immediately for DIC transmitted light and green autofluorescence (GAF, excitation 488 nm, emission 495 to 550 nm). The cell culture plate was placed in the dark with gentle shaking. **(B)** 11 hours after subjective dusk, cells had divided, and the second coverglass was imaged. The media had regained a green color, presumably from released autospores no longer bound to the coverglass. Putative shed red bodies and autosporangial walls remained affixed. **(C)** The green colored media was transferred to a new well, and cells re-bound onto a new coverglass and imaged. These recently divided cells are smaller and have not yet developed visible red bodies. The scale bar = 5 μm in all images.

The cell wall of *Nannochloropsis* has been found to be made of two main layers, a thick cellulose inner layer, and a thinner, outer layer made of a hydrophobic, chemically recalcitrant polymer called algaenan (Scholz et al., 2014). In the late stages of cell division, we directly visualized the red body outside of developing autospore cell walls, but enclosed by the enveloping autosporangial wall by labeling cellulose with the fluorescent dye, calcofluor white (CFW) (Figure 4, supplemental 1). We additionally observed shed red bodies within CFW-labeled shed autosporangial walls, which suggests that these shed walls retain at least some cellulose from the inner cell wall layer (Figure 4, supplemental 1).

### Cells exhibit a period of increased permeability to exogenous dyes that coincides with autospore release from the enclosing autosporangial wall

In the CFW staining experiment above, relatively few cells appeared to be labeled. Low efficiency in stain uptake has been reported for this species and close relatives (Doan and Obbard, 2011; Südfeld et al., 2021; Veldhuis et al., 1997), and in other algae that produce algaenan (Dunker and Wilhelm, 2018; Zych et al., 2009). Algaenan cell coverings may serve as protection for resting spores against desiccation (Goodenough et al., 2007; Heimerl et al., 2018) or as defense against biotic stress (Dunker and Wilhelm, 2018), and it may thus form a physical barrier to exogenous dye entry.

We hypothesized that there may exist a window of increased permeability as the autosporangial wall degrades in preparation for daughter cell release, but before the new autospore cell walls have fully matured. We thus incubated synchronously dividing cells with CFW or the DNA-binding dye, Hoechst 33342, for 3-hour periods starting at different points during subjective night, washing cell aliquots to reduce background unbound dye, and observing fluorescence of these cells in bulk culture and by microscopy (schematic in Figure 5A; synchrony check in Figure 5, supplemental 1). Cells incubated with the dyes during the period of autospore release (dusk +3 to +7 hours) exhibited greater rates of fluorescence increase (dye uptake) compared with time windows a few hours before or after (Figure 5B, C). That is, the slopes of a line of best fit were different between windows (Figure 5B). An analysis of covariance (ANCOVA) with the model Fluorescence ∼ Time * Window yielded significant interaction effects between time and staining window for the CFW treatment (p-value = 1.66e-7), Hoechst (p-value = 1.44e-12), but not for the no-stain control (p-value = 0.245). This indicates that the slopes (rate of dye uptake or staining frequency) are different between the staining windows.

**Figure 5.**
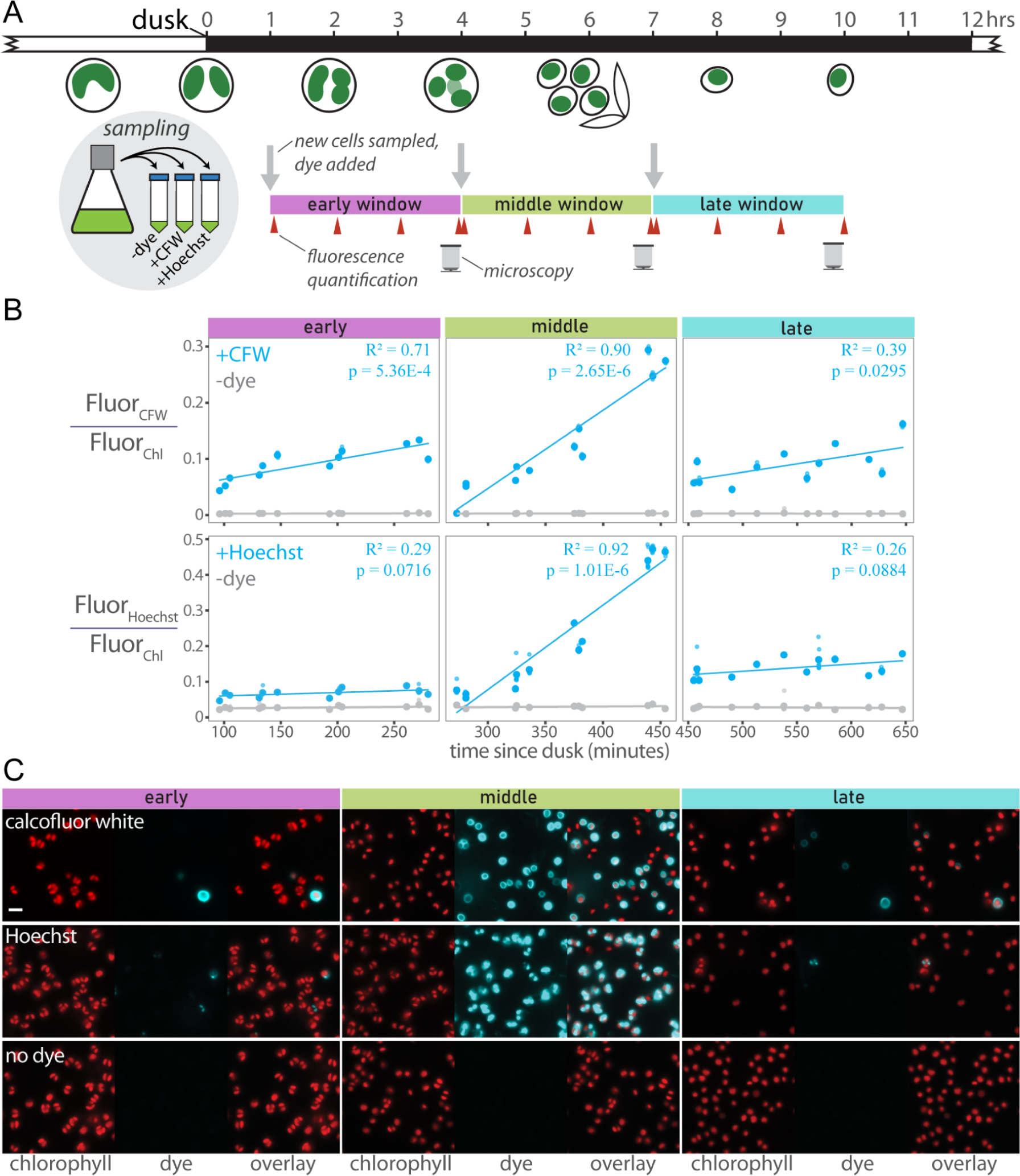
Cells exhibit increased permeability to exogenous dyes during the period of autospore maturation and release. **(A)** Synchronized cells were sampled from a culture at the indicated times after dusk. Sampled culture was divided into three treatments: no dye addition, calcofluor white (CFW), or Hoechst 33342. These treatments were subsampled every hour, washed of excess dye, and fluorescence measured on a plate reader (red arrowheads). At the end of the 3 hour window, washed cells were imaged; CFW and Hoechst fluorescence pseudo colored as cyan, chlorophyll fluorescence as red. These procedures were repeated with new samples from the original culture two more times. The entire experiment was carried out three times on different days. **(B)** Fluorescence over time from CFW or Hoechst was normalized to chlorophyll fluorescence. The three experimental runs were plotted together, and each data marker is the mean of 7 technical replicates (individual wells), which are represented by smaller data markers that generally lie inside the larger markers. Simple linear regression lines, coefficients of determination, and overall p-values are shown for each treatment in each panel. **(C)** Widefield fluorescence microscopy. For each window, chlorophyll fluorescence (red), dye fluorescence (cyan), and an overlay are shown. The scale bar in the upper left image = 5 μm, and applies to all images.

Microscopy of stained cells revealed circular outlines (cell walls) for the CFW-dyed cells and multiple punctae (nuclei) in each cell for the Hoechst dyed cells, consistent with their expected binding targets (Figure 5C, additional microscopy in Figure 5 supplemental 2). The frequency of staining was relatively low in the early and late windows compared to the middle window (Figure 5C, Figure 5C supplemental 2). Together, these observations are consistent with a gap in algaenan coverage between the degradation of the maternal autosporangial wall and the formation of new algaenan layers around autospores.

### Shed cell walls and red bodies contain ketocarotenoids and a long-chain alkyl diol

We observed a “red sediment” on top of cell pellets of *Nannochloropsis* when harvesting large numbers of cells by centrifugation (Figure 6A). This extracellular debris has also been reported in a study of *Nannochloropsis* media recycling (Rodolfi et al., 2003). This red sediment consisted of particles matching the appearance of shed autosporangial walls and red bodies (Figure 4, Figure 6B). TEM revealed the rolled and wrinkled appearance of the autosporangial walls (Figure 6C) that were similar in thickness (∼5 nm) to the outer, algaenan cell wall layer of *Nannochloropsis* (Figure 6D) (Scholz et al., 2014). The shed red bodies appeared as roughly spherical and homogenous particles of slightly less than ∼300 nm in diameter (Figure 6C, D).

**Figure 6.**
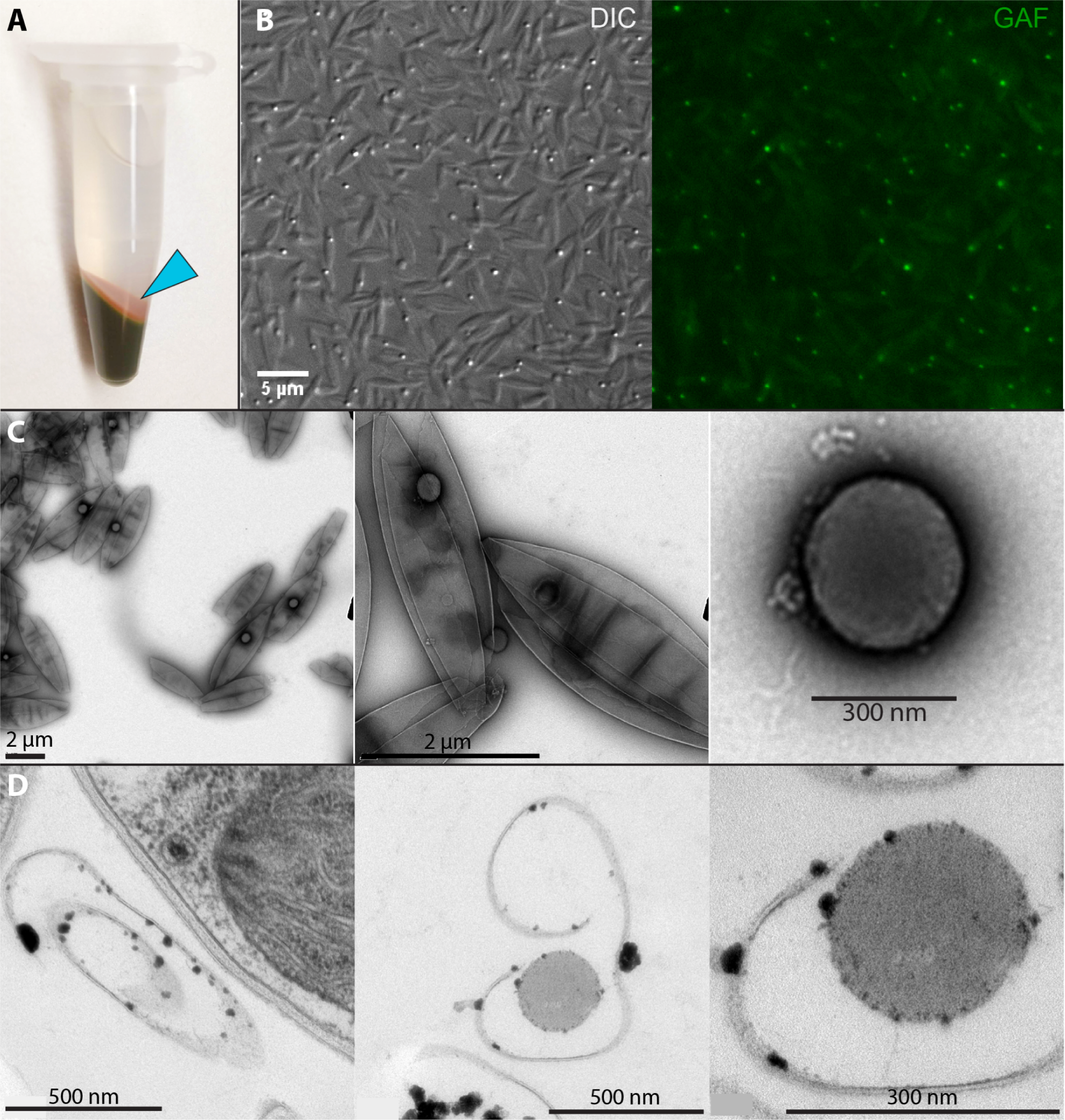
Shed cell walls and red bodies can be collected from the media. **(A)** 100 mL of culture was concentrated into 1 mL by centrifugation, showing the red sediment on top of the cell pellet (cyan arrowhead). **(B)** This sediment imaged by fluorescence microscopy shows elongated, pointed particles and smaller spherical particles that exhibit fluorescence similar to *in vivo* red bodies. DIC transmitted light and green autofluorescence (GAF) acquired as described previously. **(C)** Whole mount, negative stained, transmission electron microscopy of the red sediment. **(D)** Resin-embedded sample of concentrated culture showing a sectioned elongate particle resembling the outer algaenan layer of a nearby intact cell (left), an apparent cross-section of a shed wall and putative red body (middle), and this same particle imaged at higher magnification (right).

Daily harvesting of dense, actively growing cultures produced relatively large amounts of the red sediment, and high pressure homogenization selectively disrupted the shed autosporangial walls, allowing us to obtain enrichments of shed red bodies (Figure 7A, Figure 7, supplemental 1). Organic solvent extracts of the red sediment and red body enrichment were subjected to ultra high pressure liquid chromatography coupled to quadrupole time-of-flight mass spectrometry (UHPLC-qTOFMS). Consistent with the observed red color of the sample, this analysis indicated the presence of the ketocarotenoids astaxanthin (m/z 597.3944 [M+H]) and canthaxanthin (m/z 565.4046 [M+H]) (Figure 7B).

**Figure 7.**
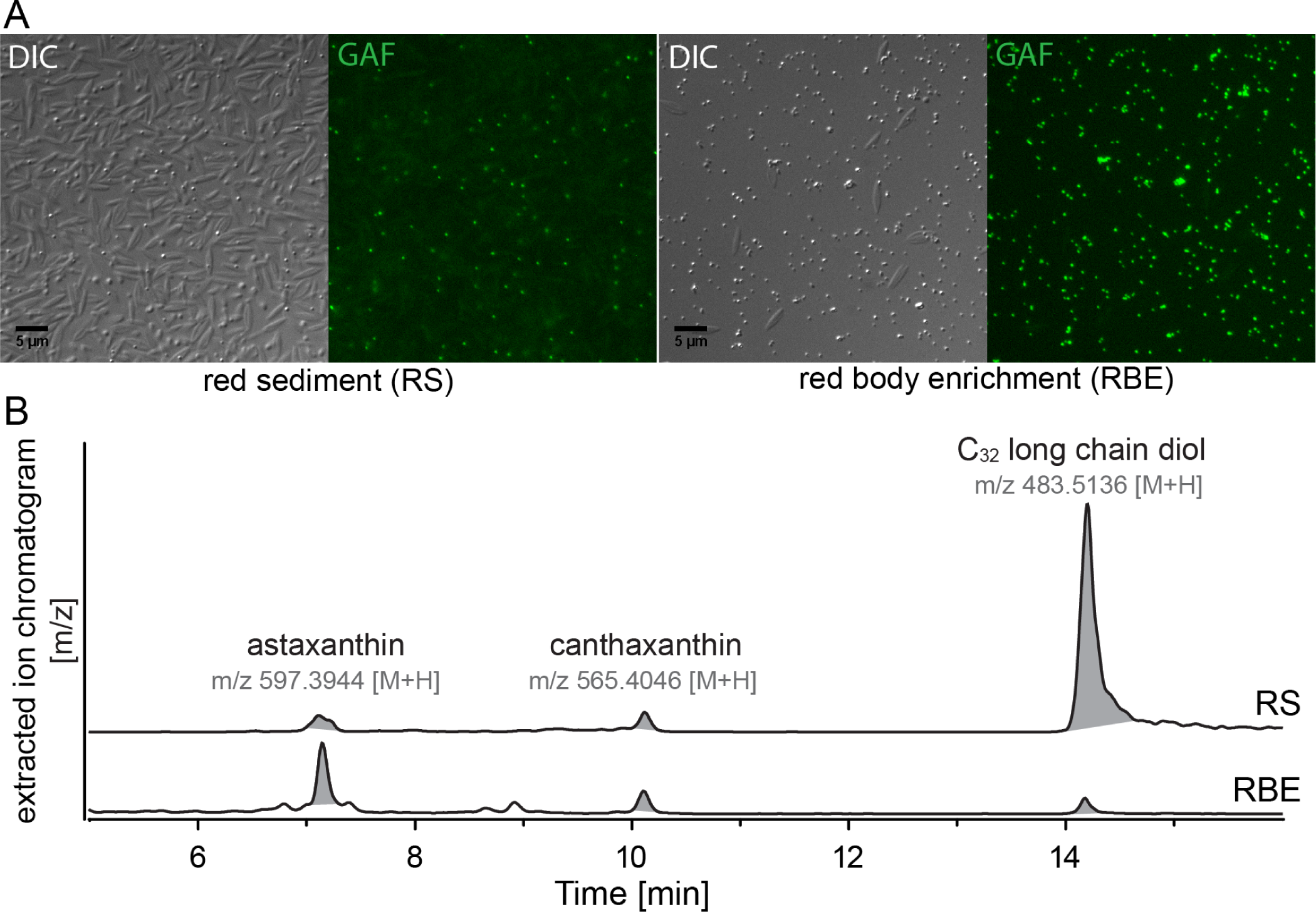
Shed cell walls and red bodies contain ketocarotenoids and a long-chain alkyl diol. **(A)** Light microscopy of the red sediment (RS) mixture and a preparation enriched in the shed red bodies (RBE). Differential interference contrast (DIC) and green autofluorescence (GAF). **(B)** Extracted ion chromatograms (EIC) from LC-HRMS in APCI mode of organic solvent extracts from red sediment (RS) and red body enrichment (RBE) samples. Shown is the EIC of m/z 597.3944, m/z 565.4046, and m/z 483.5136 ± m/z 0.01 for identification of astaxanthin, canthaxanthin, and C_32_ long-chain diols, respectively. Identity of astaxanthin and canthaxanthin was confirmed by authentic standards. Identification of the C_32_ long-chain diols was based on its accurate mass (< 5 ppm) and the observed in-source fragmentation of this compound in APCI (Figure 7-supplemental 2).

Published reports of the chemical composition of *Nannochloropsis* algaenan indicated a structure made primarily of unbranched C_30-32_ alkyl diols (terminal and mid-chain –OH) in which the hydroxyl groups have been converted to ether linkages between alkyl chains (Gelin et al., 1996; Scholz et al., 2014; Zhang and Volkman, 2017). In *Nannochloropsis*, the precursors to algaenan are thought to be long-chain diols (LCDs), as well as mono- and di-hydroxylated fatty acids that are reduced to form the LCDs (Balzano et al., 2019, 2017). In the UHPLC-qTOFMS, we identified a molecule with a parent mass of m/z 483.5117 [M+H], which corresponds to a predicted molecular formula C_32_H_66_O_2_ (mass error = −3.84 ppm) (Figure 7B), likely being the LCD dotriacontane-1,15-diol. In the mass spectra of the detected LCD, m/z 465.5012 and 447.4896 were also observed. These correspond to putative in-source fragmentation products of the LCD without one or both of the -OH groups (mass error = −3.65 ppm & −6.32 ppm, respectively) (Figure 7, supplemental 2). Relative to the observed amount of LCD, ketocarotenoids were more abundant in the red body enrichment sample, while the opposite was observed in the red sediment sample (Figure 7B). Because the red sediment contains much more shed wall material, this suggests that, for extractable molecules, the LCD originated primarily from the shed wall, and the carotenoids from the red body.

Accumulation of ketocarotenoids has been observed in several green algae including *Chromochloris zofingiensis*, in which beta-carotene ketolase (*BKT*) catalyzes the addition of the characteristic ketone groups (Roth et al., 2017; Ye and Huang, 2020). A clear homolog of green algal BKT was not found in the *N. oceanica* genome so we ectopically expressed the *C. zofingiensis BKT* (*CzBKT*ox) coding sequence, with and without fusion to an *N. oceanica* chloroplast targeting peptide (cTP) (Moog et al., 2015). Only transformation of the cTP fusion construct into *N. oceanica* produced any colonies with a noticeable brown tint, which yielded cultures with an obvious brown color (Figure 8A). Bulk cell suspensions had increased absorbance between 460 nm - 580 nm, suggesting increased carotenoid content (Figure 8B). Comparing the extracted ion chromatograms from LC-DAD-mSqTOF analysis, the *CzBKT*ox cells yielded more canthaxanthin than astaxanthin relative to wild-type cells, and the intermediate between the two, adonirubin, was only clearly detected in *CzBKT*ox cells (Figure 8C). Strikingly, we observed unusually large red bodies in these cells, as well as the presence of cells with multiple red bodies (Figure 8D). Because significant keto-carotenoid accumulation only was observed upon expression of cTP-*Cz*BKT, which directed the enzyme to the plastid where endogenous carotenoid biosynthetic enzymes are located, we interpret these observed effects on the red body as evidence of metabolic flux between the chloroplast and red body.

**Figure 8.**
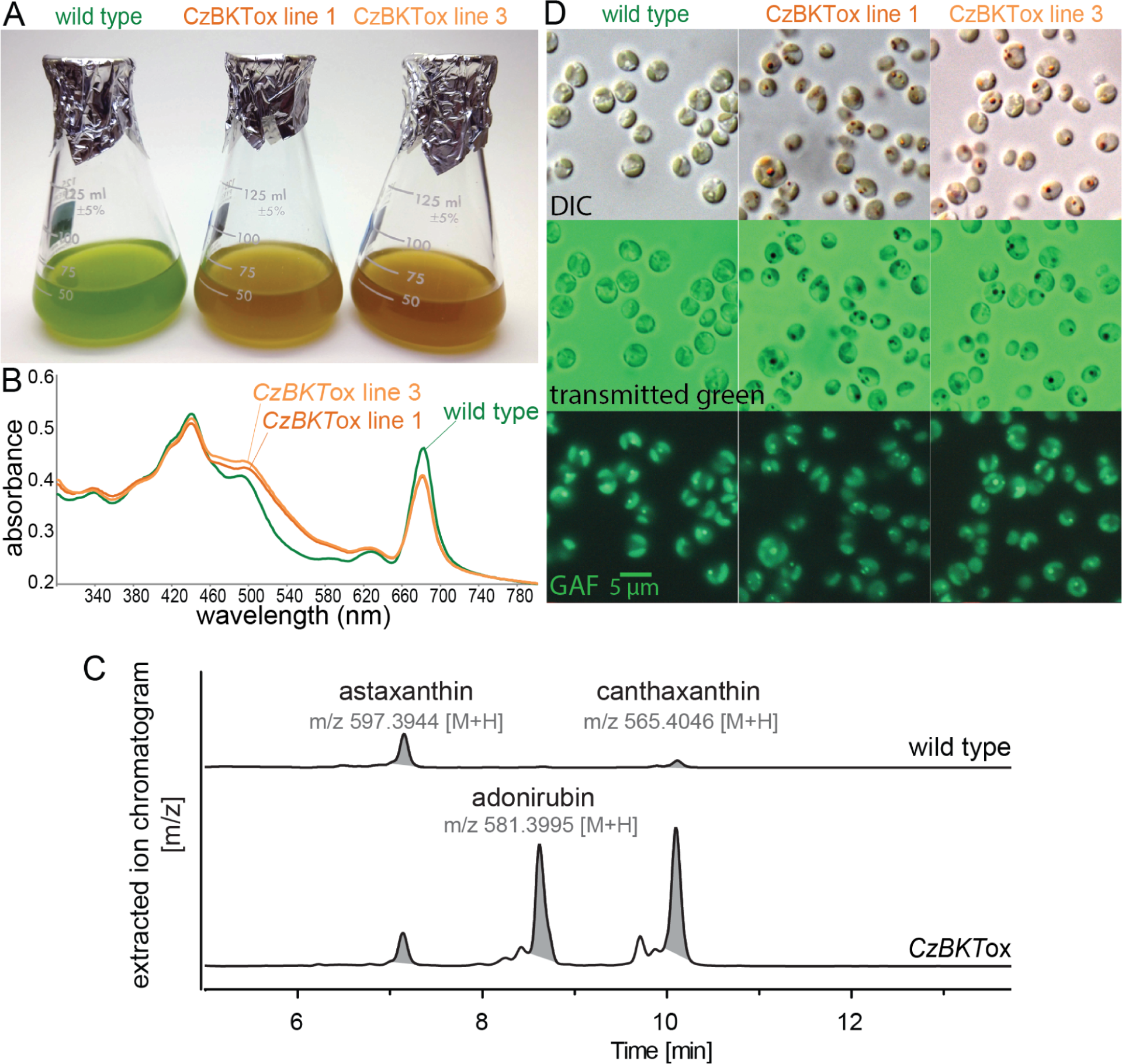
Ketocarotenoid overaccumulation leads to aberrant red bodies. **(A)** *Nannochloropsis oceanica* cultures of wild type and two independent lines overexpressing a beta carotene ketolase from the green alga, *Chromochloris zofingiensis* (*CzBKT)*. Cultures were adjusted to equal cell density for imaging. **(B)** Absorbance spectra for the same cell suspensions, normalized to OD_800_ (primarily scattering). **(C)** LC-DAD-HRMS analysis of extracts from wild type and *CzBKTox* cells. Shown is the EIC of m/z 597.3944, m/z 565.4046, and m/z 581.3995 ± m/z 0.01 for identification of astaxanthin, canthaxanthin, and adonixanthin, respectively. Astaxanthin and canthaxanthin was confirmed by authentic standards while adonixanthin identification was based on accurate mass (< 5 ppm) and comparison to reference absorbance spectra (Britton et al., 2004). **(D)** Light microscopy showing differential interference contrast (DIC), transmitted green light (broad spectrum halogen lamp viewed through the GFP filter set), and green autofluorescence (GAF). Scale bar = 5 µm and applies to all images.

### Infrared spectroscopy indicates shed cell walls and red bodies both contain saturated lipids, but differ in carbohydrate and carbonyl-associated molecules

To obtain even greater enrichment of the shed red body for infrared spectroscopy, we extended the preparation method from the pigment analysis (Figure 7, supplemental 1) with ultracentrifugation through a sucrose density gradient, followed by water washes (Figure 9, supplemental 1A and B). Additionally, we found that by incubating the red sediment mixture in 1% SDS at 50 °C, the red bodies were solubilized, leaving behind the shed autosporangial walls (Figure 9, supplemental 1A and B). Cultures of another eustigmatophyte, *Goniochloris sculpta*, also contained extracellular debris that appeared to be shed cell walls that could be isolated by differential sedimentation. However, extracellular red bodies from this species were not obvious (Figure 9, supplemental 1C). These various isolations were prepared and analyzed by attenuated total reflection infrared (ATR-IR) spectroscopy (Cui et al., 2016). Overview spectra with general annotations are shown in Figure 9 (detailed peak positions in Figure 9, supplemental 2; higher order derivative analyses in Figure 9, supplemental 3), and detailed relevant peak assignments are summarized in Table 1.

**Figure 9.**
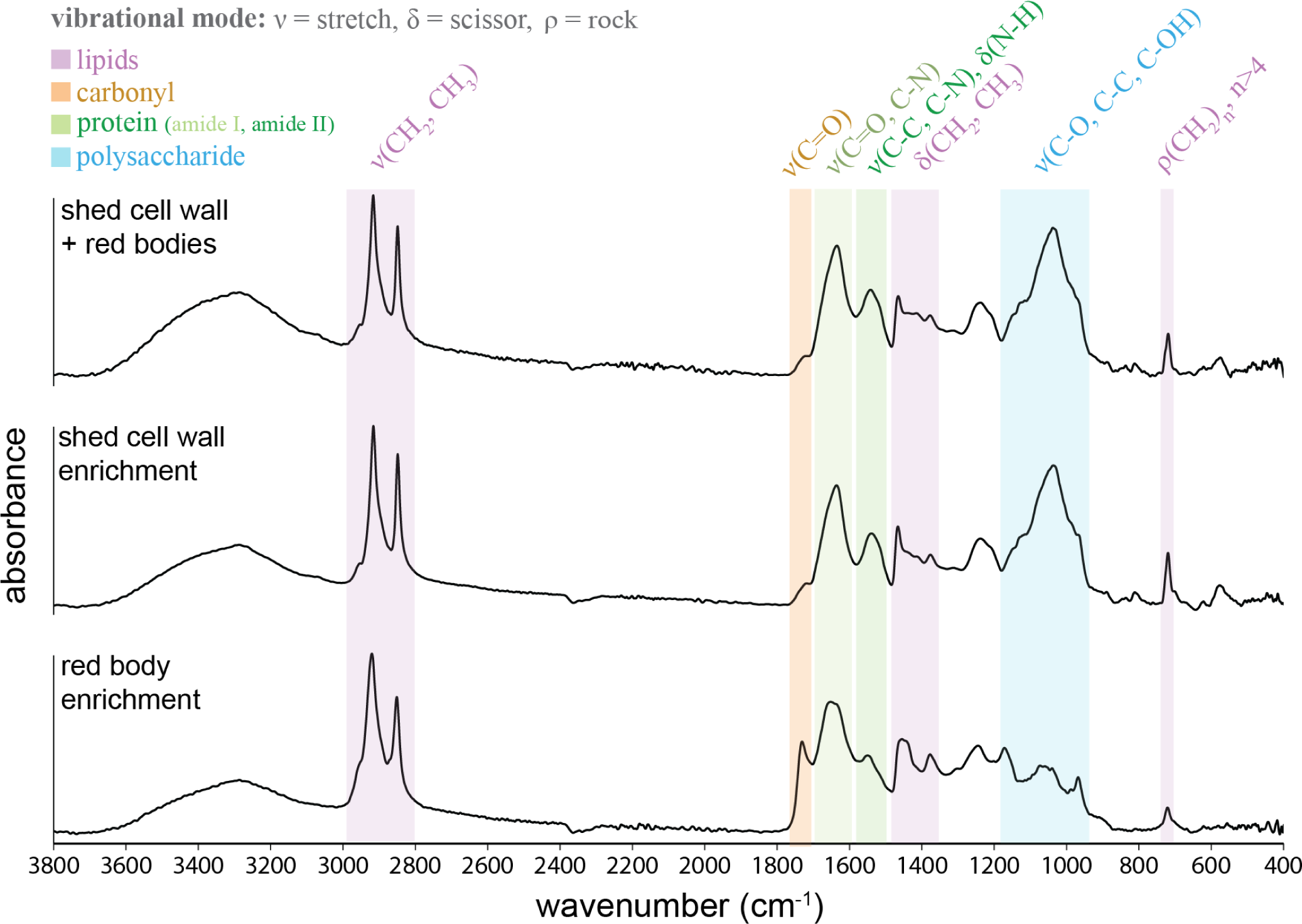
ATR-FTIR spectroscopy indicates shed cell walls and red bodies have different lipid and carbohydrate composition. Attenuated total reflectance Fourier transform infrared (ATR-FTIR) spectroscopy of shed cell walls and red bodies (red sediment) and preparations enriched in only the shed cell walls or the red bodies. Baseline corrected absorbance spectra are shown, vertically scaled to equal heights. Molecular vibration assignments are annotated above bands that are color coded with their associations to the dominant biological classes of molecules. Classes, their label colors, and approximate wavenumber locations (cm^−1^) are as follows: lipids (purple: 3100 - 2800, 1500 - 1300, 1240 and 1171, 721-718); non-peptide carbonyl (orange: 1750 - 1700), protein (light green for Amide I: 1690 - 1600, dark green for Amide II: 1600-1480), polysaccharide (blue: numerous features between 1300 - 900).

**Table 1.**
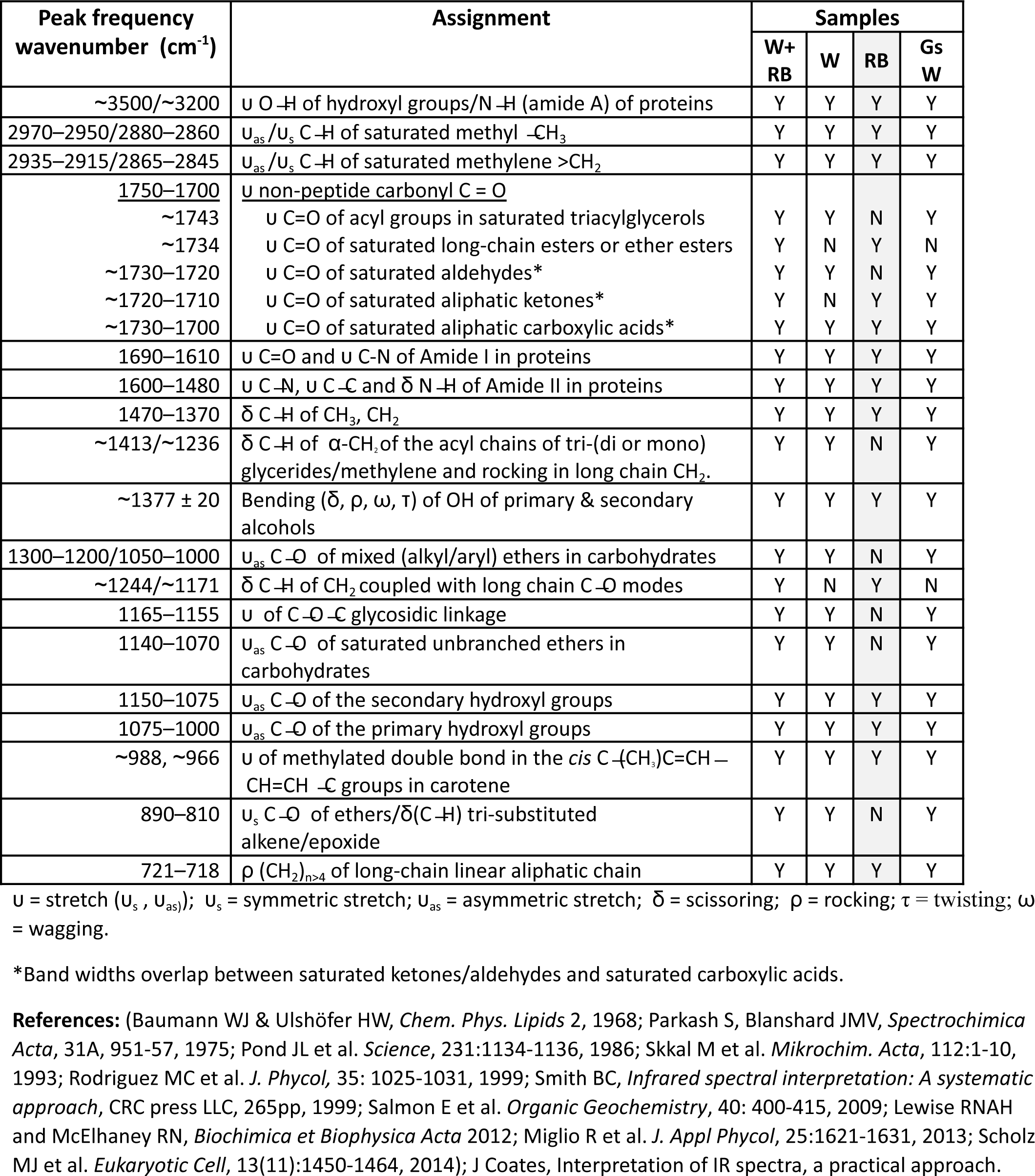
Characteristic absorption frequencies in the mid-infrared spectral range from 2000 to 650 cm^−1^ for **(W+RB)** *N. oceanica* shed cell walls and red bodies, **(W)** *N. oceanica* shed cell walls, **(RB)** *N. oceanica* shed red bodies, and **(Gs W)** *G. sculpta* shed cell walls.

| Peak frequency wavenumber (cm<sup>-1</sup>) | Assignment | Samples |  |  |  |
| --- | --- | --- | --- | --- | --- |
|  |  | W+RB | W | RB | Gs W |
| ~3500/~3200 | υ O-H of hydroxyl groups/N-H (amide A) of proteins | Y | Y | Y | Y |
| 2970–2950/2880–2860 | υ<sub>as</sub>/υ<sub>s</sub> C-H of saturated methyl -CH<sub>3</sub> | Y | Y | Y | Y |
| 2935–2915/2865–2845 | υ<sub>as</sub>/υ<sub>s</sub> C-H of saturated methylene >CH<sub>2</sub> | Y | Y | Y | Y |
| 1750–1700 | υ non-peptide carbonyl C=O |  |  |  |  |
| ~1743 | υ C=O of acyl groups in saturated triacylglycerols | Y | Y | N | Y |
| ~1734 | υ C=O of saturated long-chain esters or ether esters | Y | N | Y | N |
| ~1730–1720 | υ C=O of saturated aldehydes* | Y | Y | N | Y |
| ~1720–1710 | υ C=O of saturated aliphatic ketones* | Y | N | Y | Y |
| ~1730–1700 | υ C=O of saturated aliphatic carboxylic acids* | Y | Y | Y | Y |
| 1690–1610 | υ C=O and υ C-N of Amide I in proteins | Y | Y | Y | Y |
| 1600–1480 | υ C-N, υ C-C and δ N-H of Amide II in proteins | Y | Y | Y | Y |
| 1470–1370 | δ C-H of CH<sub>3</sub>, CH<sub>2</sub> | Y | Y | Y | Y |
| ~1413/~1236 | δ C-H of α-CH<sub>2</sub> of the acyl chains of tri-(di or mono) glycerides/methylene and rocking in long chain CH<sub>2</sub>. | Y | Y | N | Y |
| ~1377 ± 20 | Bending (δ, ρ, ω, τ) of OH of primary & secondary alcohols | Y | Y | Y | Y |
| 1300–1200/1050–1000 | υ<sub>as</sub> C-O of mixed (alkyl/aryl) ethers in carbohydrates | Y | Y | N | Y |
| ~1244/~1171 | δ C-H of CH<sub>2</sub> coupled with long chain C-O modes | Y | N | Y | N |
| 1165–1155 | υ of C-O-C glycosidic linkage | Y | Y | N | Y |
| 1140–1070 | υ<sub>as</sub> C-O of saturated unbranched ethers in carbohydrates | Y | Y | N | Y |
| 1150–1075 | υ<sub>as</sub> C-O of the secondary hydroxyl groups | Y | Y | Y | Y |
| 1075–1000 | υ<sub>as</sub> C-O of the primary hydroxyl groups | Y | Y | Y | Y |
| ~988, ~966 | υ of methylated double bond in the *cis* C-(CH<sub>3</sub>)C=CH-CH=CH-C groups in carotene | Y | Y | Y | Y |
| 890–810 | υ<sub>s</sub> C-O of ethers/δ(C-H) tri-substituted alkene/epoxide | Y | Y | N | Y |
| 721–718 | ρ (CH<sub>2</sub>)<sub>n>4</sub> of long-chain linear aliphatic chain | Y | Y | Y | Y |

υ = stretch (υ<sub>s</sub>, υ<sub>as</sub>); υ<sub>s</sub> = symmetric stretch; υ<sub>as</sub> = asymmetric stretch; δ = scissoring; ρ = rocking; τ = twisting; ω = wagging.
\*Band widths overlap between saturated ketones/aldehydes and saturated carboxylic acids.
**References:** (Baumann WJ & Ulshöfer HW, *Chem. Phys. Lipids* 2, 1968; Parkash S, Blanshard JMV, *Spectrochimica Acta*, 31A, 951-57, 1975; Pond JL et al. *Science*, 231:1134-1136, 1986; Skkal M et al. *Mikrochim. Acta*, 112:1-10, 1993; Rodriguez MC et al. *J. Phycol*, 35: 1025-1031, 1999; Smith BC, *Infrared spectral interpretation: A systematic approach*, CRC press LLC, 265pp, 1999; Salmon E et al. *Organic Geochemistry*, 40: 400-415, 2009; Lewis RNAH and McElhaney RN, *Biochimica et Biophysica Acta* 2012; Miglio R et al. *J. Appl Phycol*, 25:1621-1631, 2013; Scholz MJ et al. *Eukaryotic Cell*, 13(11):1450-1464, 2014); J Coates, *Interpretation of IR spectra, a practical approach*.

All spectra contained very notable absorbance peaks associated with unbranched, long-chain, saturated lipids. These include a strong pair at ∼2916 cm^−1^ and ∼2848 cm^−1^ (anti-symmetric and symmetric C ̶ H stretching modes of methylene groups, CH_2_) with a small shoulder at ∼2954 cm^−1^ (anti-symmetric C ̶ H stretching mode of terminal methyl groups, CH_3_), a moderate pair near 1460 cm^−1^ and 1375 cm^−1^ (CH_2_ and CH_3_ scissoring), and a peak near 720 cm^−1^ (rocking of more than 4 adjacent methylene) (Figure 9, purple bands) (Larkin, 2011; Lewis and McElhaney, 2013). The higher absorption intensity and a red-shift of the methylene rocking band from ∼721 cm^−1^ in the red body to ∼718 cm^−1^ in the shed wall sample indicate a longer alkane chain length in the shed wall.

IR spectra for both red bodies and shed walls showed high-intensity protein bands at 1700 – 1600 cm^−1^ (C=O stretch in amide I groups) (Figure 9, light green band) and 1580 – 1480 cm^−1^ (N ̶H/C ̶ N stretch in amide II groups) (Figure 9, dark green bands). The shift of the max of the amide I band from ∼1648 cm^−1^ in the red body to ∼1635 cm^−1^ in the cell wall implied a change of the dominant protein secondary structure from unordered in the red body to β-sheet in the cell wall (Barth, 2007).

The spectra differed notably in the non-peptide carbonyl region (1750-1700 cm^−1^; C=O stretch) (Figure 9, orange band). The spectra from red bodies contained a prominent peak at ∼1731 cm^−1^ not found in other samples, and further analyses suggested the presence of long-chain fatty acids, esters / ether esters (Baumann and Ulshöfer, 1968; Jones, 1962; Parkash and Blanshard, 1975; Rodriguez et al., 1999). The peak in the raw spectrum at ∼1731 cm^−1^ arose from two overlapping subcomponents at ∼1734 cm^−1^ (carbonyl stretch in saturated long-chain esters or ether esters) and ∼1710 cm^−1^ (carbonyl stretch in saturated carboxylic acids or ketones) (Figure 9, supplemental 3A). The presence of esters and ether ester functional groups was supported by the strong pair at ∼1244 cm^−1^ and ∼1171 cm^−1^ (C ̶ H deformation of CH_2_ groups of the long-chain moiety coupled to the C ̶ O vibrations of C ̶ O ̶ C) (Figure 9, supplemental 2). For shed walls, the strong carbonyl shoulder at ∼1718 cm^−1^ is a summation of two overlapping components with maxima at ∼1743 cm^−1^ (carbonyl stretch in acyl groups of saturated triglyceride moiety) and ∼1722 cm^−1^ (carbonyl stretch in aliphatic aldehydes or carboxylic acids) (Figure 9, supplemental 2 and 3A). As a consistency check, the subcomponent peaks found in only the samples of shed walls, or only red bodies were all present together in the 2^nd^ derivative spectrum of the red sediment mixture of the two (Figure 9, supplemental 3A).

Shed red bodies further differed from shed cell walls in polysaccharide structure and abundance. Shed walls exhibited a prominent, broad peak centered at ∼1038 cm^−1^ (C ̶ O ̶ C stretch of alkyl ether) in the region of 1140-1000 cm^−1^ (common to cellulose molecules due to coupling of the C ̶ O / C ̶ C stretch with the C ̶ OH bending or C ̶ O ̶ C ether bridge (Higgins et al., 1961) (Figure 9, blue band). This typical spectral feature of cellulose was missing in the red body enrichment spectrum. In mechanically isolated cell walls of whole *Microchloropsis gaditana* cells, a similar polysaccharide-associated feature was shown to be greatly diminished by cellulase treatment (Scholz et al., 2014). These authors also assigned a narrow band at ∼1157 cm^−1^ to the glycosidic C ̶ O ̶ C ether stretch, which we also found in higher derivative analyses of the shed wall enriched sample spectrum (Figure 9, supplemental 3C). Altogether, these data strongly suggest the presence of algal cellulose in the shed cell wall and relative scarcity in shed red bodies.

Despite belonging to an entirely different order-level clade within the class Eustigmatophyceae relative to *Nannochloropsis* (Figure 1; (Amaral et al., 2020)), the infrared spectra of shed cell walls of *Goniochloris sculpta* generally resembled those of *N. oceanica*. The peaks from *Nannochloropsis* indicating long-chain, unbranched lipids (e.g. ∼2909 cm^−1^ and ∼2850 cm^−1^, ∼1463 cm^−1^, ∼1375 cm^−1^ and ∼718 cm^−1^), amide bands (∼1634 cm^−1^ and ∼1530 cm^−1^), and cellulose (broad peak at ∼1043 cm^−1^, narrow peak at ∼1159 cm^−1^) were present in the *G. sculpta* cell walls, although with different exact locations (Figure 9, supplemental 2 and 3). Of course, in addition to the broad similarities, there were many differences in exact location and entirely present / absent peaks between the two species (Figure 9, supplemental 2 and 3).

The globular form and carotenoid content of the red body suggested similarities between this organelle and pigmented triglyceride storage bodies in other algae (Ota et al., 2018; Rabbani et al., 1998; Wang et al., 2019). However, the non-peptide carbonyl peak position from shed red bodies is not consistent with triglycerides (Figure 9, supplemental 3). Further, we isolated shed red bodies and subjected them to ultracentrifugation though discontinuous sucrose gradients to roughly estimate their buoyant density. These isolated red bodies passed through 24% but not 30% sucrose solution (weight/volume at 20°C), bounding their density between 1.103 - 1.127 g/cm^3^ (Figure 9, supplemental 4) (Anderson, 1966). This is considerably denser than would be expected for typical lipid bodies, which sediment on top of aqueous centrifugation layers (0.998 g/cm^3^) (Ding et al., 2013).

### Hypothesis: The red body is an organelle that carries algaenan precursors to the apoplast for cell wall biosynthesis during cell division

Our combined observations led us to the hypothesis that the red body is an organelle that carries material into the apoplast during the late stages of cell division and autospore development. These molecules may include plastid-derived lipid precursors of the algaenan polymer and proteins required for its biochemical maturation. The physical and metabolic connections with the plastid, the secretion into the apoplast within the enclosing autosporangial wall, and the presence of long-chain, saturated lipids (matching expected composition of algaenan precursors) are consistent with this proposed function.

### Red body organelle volume is correlated with cell size and declines during autospore maturation

If the red body accumulates algaenan precursors, we expected that larger parental cells would have larger red bodies, as the resulting autospores would be larger, and require a greater amount of new cell wall material to match the increase in cell surface area. Secondly, we expected that the red body would decrease in volume as material is released during the deposition of autospore cell walls.

By growing cells at elevated CO_2_ (3%) under three different light intensities (20, 60, and 180 μmol photons m^−2^ s^−1^), we were able to obtain cells of different maximum size (Figure 10A). Maximum red body volume, as estimated from super-resolution fluorescence microscopy, occurred around subjective dusk, and appeared to increase with cell size (Figure 10B, C). Cell division was completed by dusk +4 hours (using chloroplast division as a proxy, Figure 10C), and this was distinct from autospore release occurring by +6 hours (Figure 10A). Red body volume decreased after subjective dusk in cells grown at any light intensity (Figure 10B). Interestingly, at the highest light level, eight autospores per initial cell were frequently observed, compared to the four typically seen in our standard 60 μmol photons m^−2^ s^−1^ condition (Figure 10C). The central tendency can be informally assessed graphically by the 95% confidence intervals for the median, given by the notches in the box plots (Figure 10B). The results of an ANOVA and post-hoc comparison test are included in Figure 10, source data 1, and the compiled measurements themselves in Figure 10, source data 2.

**Figure 10.**
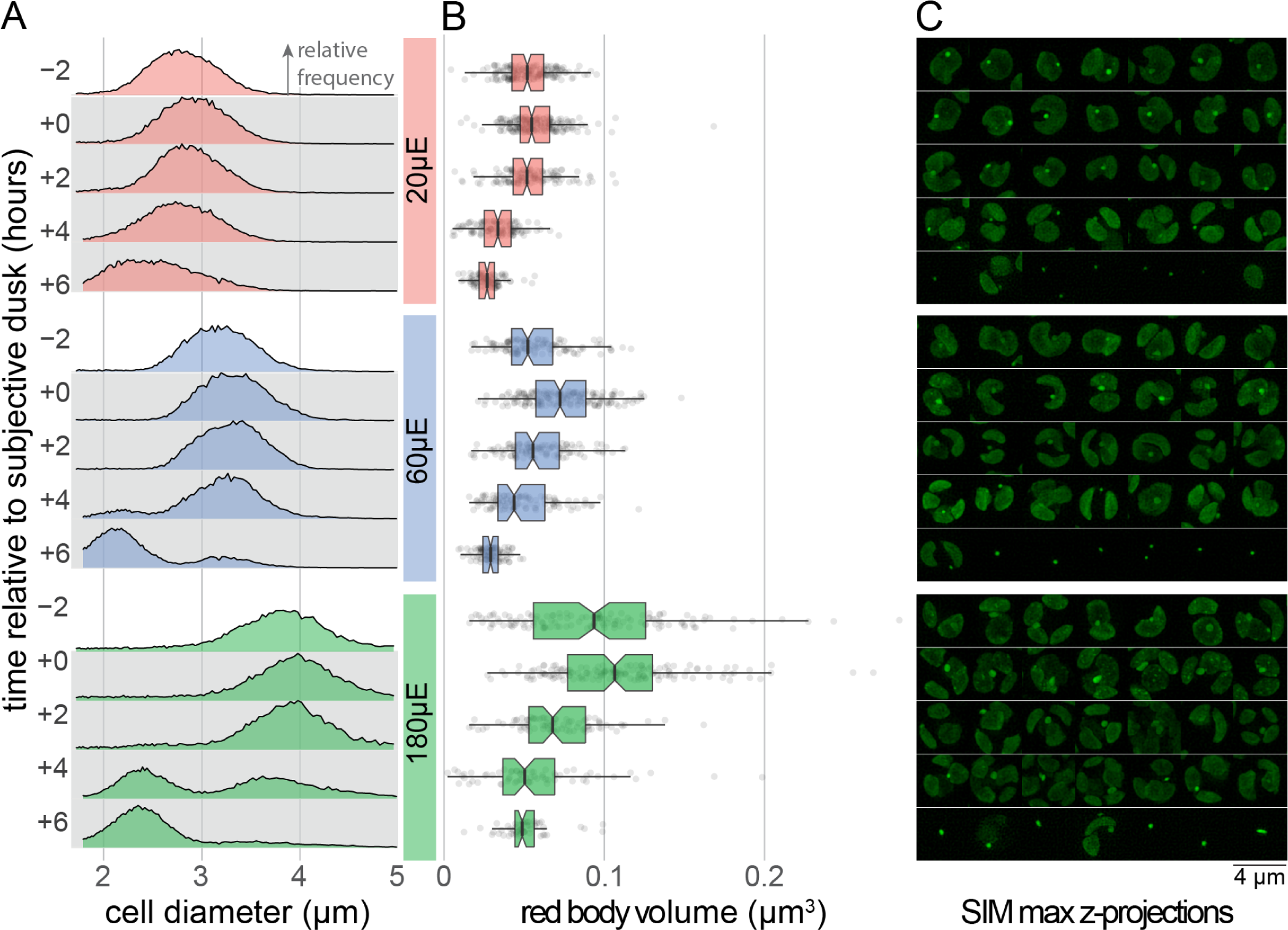
Red body volume is correlated with cell size, and it declines during autospore maturation. **(A)** Synchronous cultures grown under 3% CO2 at light levels of 20, 60, and 180 μmol m^−2^ s^−1^ (abbreviated as “μE”). Cell size was quantified by Coulter counter at the stated timepoints relative to subjective dusk, and the resulting distributions are shown as relative frequency. Bin size ∼0.01 μm, maximal y-axis height for whole panel ∼3.5%, n (counts) per distribution >13k. **(B)** Volumetric estimates derived from SIM z-stacks of cells sampled at the same timepoints as in (A). Notched box plot shows median and quartiles, with individual data points plotted behind. The notch width corresponds to the 95% confidence interval for the location of the median. ANOVA and post-hoc test results for all comparisons included in Figure 10- source data 1. The n for each distribution was between 56 and 204 (exact n included in Figure 10- source data 2). **(C)** SIM maximum intensity z-projections of randomly-selected example cells. All panels share the same time-related y-axes.

## Discussion

### The algaenan biogenesis hypothesis for red body form and function

We propose that the red body of *N. oceanica* functions in algaenan biosynthesis by compartmentalizing precursor molecules along with ancillary proteins and small molecules as they are synthesized during the daily growth period. Later, during cell division at night, the red body then delivers these molecules to the apoplast where they generate the outer algaenan layer of autospore cell walls (Figure 11).

**Figure 11.**
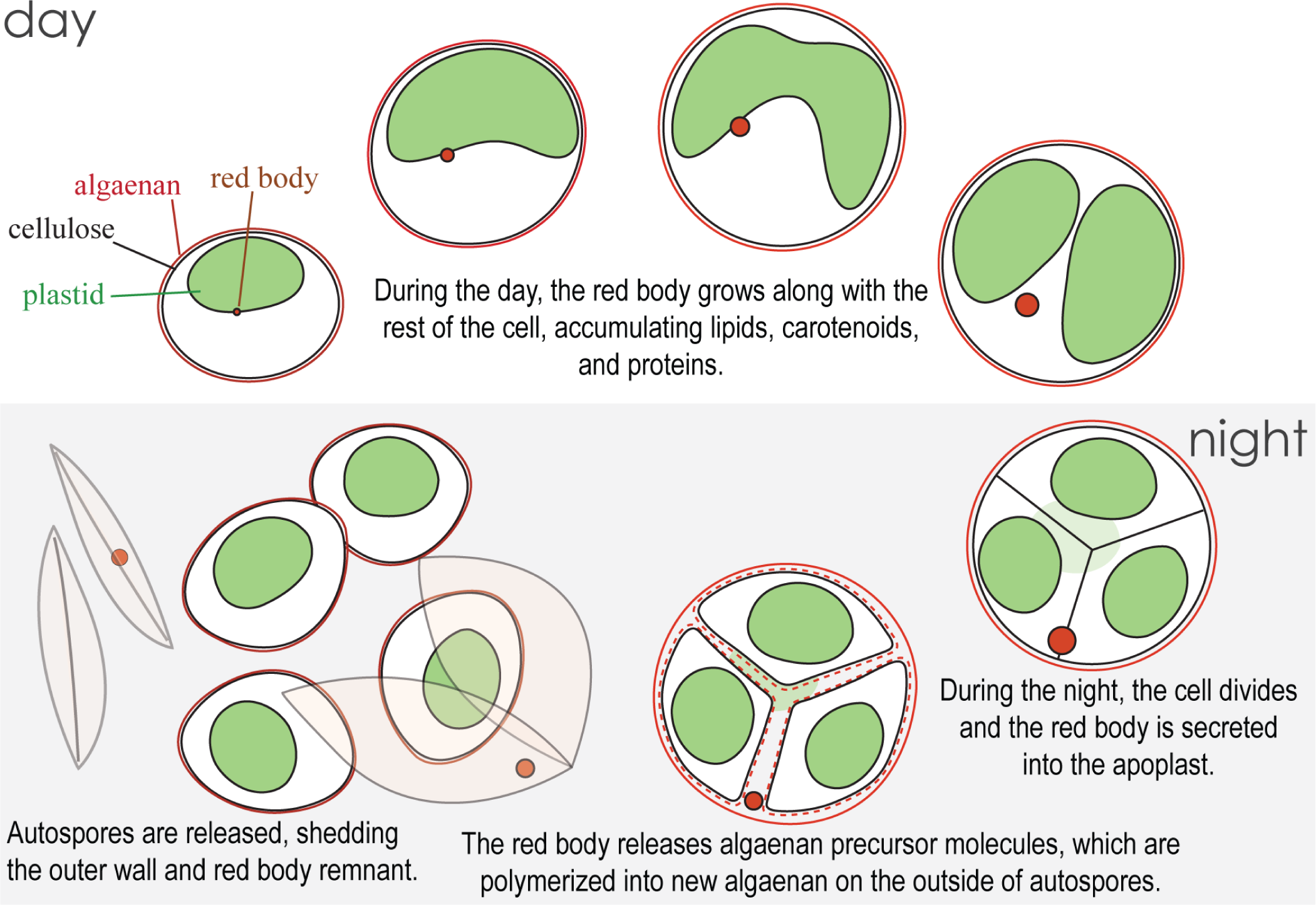
A proposed model for the development and function of the red body in *Nannochloropsis oceanica*.

Our chemical analyses support the presence of algaenan precursors in the shed red bodies and provide intriguing connections to previous work on algaenan. *Nannochloropsis* algaenan is thought to be composed of long (mostly C_32_), straight, saturated alkyl chains connected by ether cross links, and the immediate precursors are proposed to be LCDs (e.g. C_32_ 1,15-diol) (Gelin et al., 1997; Scholz et al., 2014; Zhang and Volkman, 2017). These studies inferred the identity of the precursors from the structure of the polymer, and here we directly detected the C_32_ 1,15-diol by parent ion and fragment mass in an organic solvent extraction of shed autosporangial walls, suggesting that there exists some free or loosely associated LCD in these shed walls (Figure 7). Notably, these ion masses were only detected at low levels in the shed red bodies themselves.

However, our infrared spectroscopic experiments showed that the shed red bodies contained methylene-rich signs expected from long, straight alkyl chains, and also a clear enrichment for non-peptide carbonyl groups (Figure 9). These carbonyls likely belong to saturated esters, carboxylic acids, or ketones (Figure 9, supplemental 3), which could represent algaenan intermediates or precursors containing more reactive functional groups. In fact, recent work tracing the biosynthetic pathway to algaenan has implicated long-chain hydroxyl fatty acids (LCHAs) as possible precursors to LCDs, and that the LCHAs themselves are derived from the condensation or elongation of saturated fatty acids (Balzano et al., 2019, 2017). These authors also identified a potential wax ester synthase that may catalyze the formation of esters from alcohols and fatty acids (Balzano et al., 2019). Thus, the infrared spectra of shed red bodies are consistent with the presence of precursors to the LCDs themselves or with intermediate products between LCDs and fully polymerized algaenan.

The localization of the red body is consistent with a role in gathering and dispersing algaenan precursors. The red body is membrane-enclosed (Figure 2, Video 1) and forms in close association with the chloroplast (Figure 3), likely enveloped by the chloroplastic ER (Figure 2). This position would facilitate the influx of the plastid-derived carotenoids that we detected (Figure 7, Figure 8) (Varela et al., 2015), as well as the long-chain alcohols, diols or fatty acid precursors that likely also originate in the plastid (Balzano et al., 2019, 2017). Formation in the chloroplastic ER would also facilitate lipid modifications to the precursors (Alboresi et al., 2016) and incorporation of proteins targeted to the secretory pathway (Moog et al., 2015). Indeed, our infrared spectroscopy revealed the presence of proteins in the shed red bodies (Figure 9), which possibly function in carrying or polymerizing algaenan precursors or in forming a core embedded component of the cell wall (Scholz et al., 2014). After its secretion (Figure 4), the now-apoplastic red body would be positioned to release algaenan precursors and proteins necessary for polymerization.

The temporal patterns of the red body “lifecycle” in *N. oceanica* are consistent with a role in cell wall biogenesis. The red body is generated *de novo* each day in healthy, growing cells, and it appears to be secreted around the time of cell division, a few hours prior to autospore release from the autosporangial wall (Figure 3). This is also the time when cells exhibited increased permeability to exogenous dyes (Figure 5), which supports the existence of a transition period from protection by the autosporangial wall to coverage by maturing autospore algaenan. In fact, a recent report utilized this period of permeability and showed an approximately 5-fold increase in transformation rate when synchronized *N. oceanica* cells were transformed during G2/M phase, compared to unsynchronized cells (Poveda-Huertes et al., 2023). Additionally, our hypothesis predicts that the red body would decrease in size during autospore maturation as precursor molecules were released. We informally observed this in 24-hour timecourse observations (Figure 3), and again when we quantified the red body volume with greater resolution (Figure 10).

An alternative hypothesis for the function of the red body could be that it serves as an eyespot like those commonly found in other algae. For example, in *Chlamydomonas reinhardtii*, carotenoid-rich bodies serve to directionally shield photoreceptors in order to orient phototaxis (Ueki et al., 2016). However, this does not explain the red body’s function in non-motile *Nannochloropsis* cells or why it would be secreted upon cell division. Secretion of cellular waste has not been extensively examined in *Nannochloropsis*, though media recycling studies indicate the presence of secreted inhibitory molecules (Rodolfi et al., 2003). The red body could function in secretion of waste products, however, the chemical similarities between the shed red body remnant and algaenan (Figure 9) and the observation that the red body decreases in size while still associated with the cell (Figure 10) are not obviously explained by this waste removal hypothesis.

### Expanding our understanding of the red body and algaenan biosynthesis

Numerous questions still surround the details of the red body biogenesis and function. What drives red body nucleation, and how is cargo targeted to the growing red body? Liquid-liquid phase separation is recognized as an important driver in organizing biological condensates (Boeynaems et al., 2018), and extending this biophysical perspective from studies of other types of lipid droplets (Thiam and Ikonen, 2021) to the red body may provide clues. The functional advantage of secreting a single, relatively large organelle instead of smaller exosomes is not clear. In fact, inheritance of the red body has been observed for some eustigmatophytes (Gao et al., 2019), so this may not be a ubiquitous strategy. While the presence of solvent-extractable long-chain diol from shed autosporangial walls suggests that algaenan precursors may be in excess, how the cell wall expands to accommodate growth while the red body is intracellular during the day period and why a remnant red body remains after autospore release are open questions.

Much remains to be determined regarding the composition of the red body and algaenan. The infrared spectroscopy of shed red bodies presented here is consistent with the presence of algaenan precursors or intermediates. Isolating and characterizing specific molecules from shed red bodies, and red bodies isolated from intact cells could take this a step further, and reveal the series of precursors leading to algaenan. Similarly, characterizing the proteins found in the red body (as also indicated by our infrared spectroscopy) could provide clues to the enzymes required for algaenan biosynthesis.

Regarding the carotenoid content of the red body, these lipophilic antioxidants might prevent oxidation and degradation of the algaenan precursors, or perhaps they could be integrated into the cell wall as structural components. Carotenoids have been found in the cell coverings of several organisms (Burczyk et al., 1981; Vermaas et al., 2008), including the rapidly growing green alga *Picochlorum celeri*, which produces an extracellular polysaccharide matrix that sediments as a red layer on top of the cell pellet (Cano et al., 2021) in a strikingly similar manner as the shed autosporangial walls and red bodies described here. Carotenoids are generally presumed to be non-fluorescent (Wolf and Stevens, 1967), though examples to the contrary can be found (An et al., 2000; Davis et al., 2014; Kleinegris et al., 2010; Vermaas et al., 2008), and so determining the biophysical mechanism underlying the apparent autofluorescence of the red body may provide insight into carotenoid fluorescence generally.

### Implications for research and applications related to *Nannochloropsis*

The existence of the eustigmatophyte red body may not be known to many researchers, which may influence interpretation of results, particularly with regards to microscopy. As examples, a hyperspectral imaging study describing carotenoid-rich lipid bodies (Davis et al., 2014) and a study using the fluorescent dye BODIPY to visualize lipid droplet accumulation (Benito et al., 2015) in *Nannochloropsis* may have misidentified the autofluorescent red bodies as triglyceride droplets.

The low staining efficiency of cells enclosed by an algaenan cell wall has been noted by other authors (Doan and Obbard, 2011; Südfeld et al., 2021; Veldhuis et al., 1997), and fixation of cells is often required to allow staining (Hyka et al., 2013). Here, we identified a window of greater permeability preceding autospore release that could prove to be useful for technical reasons. This knowledge also might lead to re-interpretation of staining experiments, as the rare cells that stain might be structurally compromised or dead.

According to recent phylogenetic analyses (Amaral et al., 2020), Figure 1 represents species from the major clades within Eustigmatophyceae. The occurrence of the red body in these different taxa from many different environments implies that the red body is an evolutionarily ancient characteristic, and the similarities in infrared spectra of *Nannochloropsis* and *Goniochloris* shed walls (Figure 8) indicate that algaenan may also be a shared, derived feature. Further work with the lesser studied eustigmatophytes will illuminate how this common organelle has diversified in form and function across the clade.

### Eustigmatophytes as model organisms to study the biosynthesis of chemically recalcitrant lipidic biopolymers

Hydrophobic, lipidic biopolymers like cutin, cutan, sporopollenin, and suberin play key roles in plant physiology by regulating the movement of water within and around these organisms. The final chemical composition varies between these polymers, however they share the same upstream, plastid-derived lipid precursors (Beisson et al., 2012; Kim and Douglas, 2013). This means that the mechanisms of their biosynthesis also share a common “problem”— how to transport hydrophobic precursor molecules through the aqueous compartments of the cell to the site of polymerization in the extracellular space. Several ABC transporters and lipid transport proteins (LTPs) have been implicated in hydrophobic polymer biosynthesis, but the details of intracellular trafficking and the final polymerization largely remain outstanding (Philippe et al., 2020; Quilichini et al., 2015).

The proposed plastid-to-apoplast transport of the red body represents one possible mechanism whereby the canonical secretory pathway may have been adapted to transport a single, large payload of cell wall components to the apoplast. The red body of *Nannochloropsis* may constitute a conceptual parallel with the orbicules of grasses, which are sporopollenin-containing granules found adjacent to maturing microspores in the anther, although the exact function of these bodies is unclear (Shi et al., 2015). Similarly, cutin monomers can coalesce into nanoparticles called cutinsomes, which may play a role in bulk transport of cutin precursors through the aqueous environment of the cell wall to the cuticle (Domínguez et al., 2015). Given our characterization of the red body, and the various advantages of *Nannochloropsis* and other eustigmatophyte algae as model organisms (rapid growth, unicellularity, genetic tractability), we propose that they may serve as promising experimental systems for future studies of lipid biopolymers and extracellular metabolism.

Lastly, better understanding hydrophobic biopolymers like algaenan at the biochemical and cellular level may improve our understanding of how they factor into biogeochemical processes at the global level. Such polymers can contribute to long-lived carbon pools, possibly due to their chemical recalcitrance to degradation (Derenne and Largeau, 2001; Gelin et al., 1999; Largeau et al., 1990; Vandenbroucke and Largeau, 2007). Increasing inputs of durable plant carbon (e.g. suberin) into soils is being explored as a climate change mitigation approach (Poirier et al., 2018). Additionally, LCDs likely originating from eustigmatophyte algae persist in freshwater sediments, and they have been studied as useful ecological and geological proxies (e.g. for temperature) (Rampen et al., 2014, 2012; Shimokawara et al., 2010). We propose that the red body is the physical context connecting LCDs (and their precursors) with algaenan cell walls, and thus it constitutes a new cell biological link in the chain between molecular and large-scale processes.

## Materials and Methods

### Strains and culture conditions

*Nannochloropsis oceanica* CCMP1779 was obtained from the Provasoli-Guillard National Center for Culture of Marine Phytoplankton (https://ncma.bigelow.org). For all experiments, media consisted of artificial seawater (final solute = 21 g L^−1^) with nutrient enrichment based on full-strength f medium (final concentrations of macronutrients: NaH_2_PO_4_ = 83 μM, NH_4_Cl = 2 mM, micronutrients as described in (Guillard and Ryther, 1962). Media was buffered by 10 mM Tris-HCl pH 8.1. Liquid cultures were shaken at ∼100 rpm in sterilized borosilicate Erlenmeyer flasks with perforated aluminum foil caps with a filter paper insert (Whatman #1, United Kingdom) for gas exchange. Small liquid cultures were maintained in polystyrene tissue culture plates (Genesee Scientific, USA). For strain maintenance, cell patches were streaked onto solid media with 0.9% bactoagar in polystyrene petri dishes.

The following strains were ordered from the SAG Culture Collection of Algae (Gottingen University, Germany): *Monodopsis unipapilla* (SAG 8.83), *Vischeria vischeri* (SAG 860-1, formerly *Eustigmatos vischeri*), *Vicheria* sp. (SAG 48.84, formerly *Chloridella neglecta*), and *Goniochloris sculpta* (SAG 29.96). It was found that Bristol media final concentration (mM): NaNO_3_ (2.94), CaCl_2_*2H_2_O (0.17) MgSO_4_*7H_2_O (0.3) K_2_HPO_4_ (0.43) KH_2_PO_4_ (1.29) NaCl (0.43)] at a pH of 8.0 worked adequately for these strains. *Goniochloris sculpta* only grew as streaks on this media with 0.8% agar. This medium was supplemented with the same trace minerals and vitamins as in *f* medium for simplicity.

Growth chambers (Percival E41L2, Perry, IA, USA) were set to 28 °C with light from fluorescent lamps (Philips Alto II F25T8) at 60 μmol photons m^−2^ s^−1^ unless otherwise stated. Generally, experiments were carried out with cells grown under ambient air (∼0.04% CO_2_), but for experiments using elevated CO_2_ conditions, 3% CO_2_ was achieved with a gas mixer and compressed CO_2_ cylinder (Alliance Gas Products, Berkeley, CA, USA). For timecourse experiments, two compartments of an algal growth chamber were each set to 12 hours light / 12 hours dark, but with offsets to accommodate reasonable sampling times (e.g. compartment 1 day period = 0:00 to 12:00, compartment 2 day period = 12:00 to 0:00).

### Cell Suspension Density and Sizing Determination

Cultures were maintained in an active growth phase (1 x 10^6^ to 1 x 10^7^ cells mL^−1^) in preparation for most experiments. Cell density and sizing was measured with a Coulter counter (Beckman Coulter Multisizer 3, CA, USA). Raw cell diameter event counts were converted to relative frequency and visualized as ridgeline plots with the R tidyverse packages (https://tidyverse.tidyverse.org/) and ggridges (https://cran.r-project.org/web/packages/ggridges/index.html). For routine culture maintenance, >1,000 counts were acquired to determine cell suspension density; for cell sizing experiments >10,000 were acquired to obtain smooth distributions.

### Light Microscopy

Prior to imaging, algal cells were adhered to microscope coverglasses, which improved efficiency and image quality by ensuring all cells in the field of view were in the same focal plane and imobile. To adhere the cells, coverglasses (#1.5 thickness, Fisher Scientific, USA) were first etched in 1 molar HCl at 50 °C for at least 4 hours, washed with copious amounts of water and dried. Prior to an imaging experiment, etched coverglasses were then placed on a hydrophobic thermoplastic sheet (Parafilm, Bemis Company Inc., WI, USA) and the top side coated with 50 mg/mL poly-D-lysine (P6407-5MG, Sigma-Aldrich, MO, USA) in 1x phosphate buffered saline for at least 1 hour at room temperature with gentle rocking. Finally, these coverglasses were rinsed in distilled water and placed in polystyrene microculture plates with algal cell or shed wall/red body suspensions, which were subsequently centrifuged at 500 rcf for 1-5 minutes to bind. These cell-coated coverslips were mounted on microscope slides and sealed with clear nail polish, and gently rinsed and dried before imaging.

Widefield fluorescence microscopy was performed with a Zeiss Axio Imager M2, with Zeiss Plan-NeoFluar 100x/1,30 oil objective (Carl Zeiss, Germany). The microscope was fitted with a QIMAGING 01-QIClick-F-M-12 monochromatic 12-bit camera (QI Imaging, CA, USA) for high-sensitivity and Zeiss MicroPublisher 5.0 RTV camera for true-color imaging. Filters used are listed here with the format “name: excitation bandpass midpoint wavelength (nm) / total bandpass width (nm), dichroic, emission bandpass midpoint / total bandpass width”. DAPI: 350/50, 400, 425 long pass. GFP: 470/40, 495, 525/50. YFP: 500/20, 515, 535/30. Cy5: 620/60, 660, 700/75. Image processing was done with the ImageJ distribution FIJI (Schindelin et al., 2012). Grayscale fluorescence images were false-colored and leveled uniformly using batch processing. DAPI images were false-colored cyan for visibility rather than a true-to-life deep blue.

A Carl Zeiss Elyra PS.1 was used for super resolution structured illumination microscope (SR-SIM). Zeiss Zen Blue software controlled acquisition with default optimized z sampling, and was additionally used for SIM processing. The red body autofluorescence was captured with a 488 nm excitation lase line and a 495-550 nm bandpass + 750 nm long pass green filter set. Chlorophyll autofluorescence was captured using a 642 nm laser line and 655 nm long pass filter. Confocal microscopy was carried out using a Carl Zeiss LSM710. Green autofluorescence from the red body was captured using a 488 nm laser line. Red chlorophyll was captured using a 633 nm excitation source.

### 3D reconstruction and volumetric estimation

SR-SIM z-stacks were analyzed with the 3D rendering and quantitation capabilities of the program Imaris (Oxford Instruments, United Kingdom). A single channel (green autofluorescence) was used for this volumetric analysis, as the red body was distinct from the plastid autofluorescence at the timepoints investigated. Segmentation was semi-automated, with consistent settings used within a light level treatment, but slight adjustments were made between light levels due to differences in size and morphology of the red bodies and were chosen to minimize any obvious disconnected, fibrous/spiky surface artifacts. These Imaris settings included background subtraction (0.1 μm, 4000 arbitrary fluorescence units), seed diameter (0.200-0.350), quality (4,000), sphericity (0.5-0.7). Following segmentation, red bodies were manually selected and their statistics exported. These .csv files were compiled and then plotted with the R ggplot2 package (note, the boxplots include a 95% confidence interval around the median with geom_boxplot (notch = TRUE). Base R functions aov() and TukeyHSD() were used for analysis.

### Resin embedded transmission electron microscopy (TEM)

Live *Nannochloropsis* cells were either centrifuged at 3000 rpm to form a pellet or were concentrated via filtration. The concentrated cell paste was placed into 2 mm wide by 50 µm or 100 µm deep aluminum freezing hats, then cryo-immobilized using a BAL-TEC HPM-010 high-pressure freezer (BAL-TEC, Inc., Carlsbad, CA). The samples were placed in freeze-substitution medium made up of 1% osmium tetroxide, 0.1% uranyl acetate, and 5% double-distilled water in acetone. All samples were freeze-substituted following the Quick Freeze Substitution method outlined in (McDONALD and Webb, 2011). Sample fragments were removed from carriers and infiltrated with Epon resin in successive steps to 100%, using centrifugation and rocking to facilitate exchange (McDonald and Müller-Reichert, 2002). Resin blocks with sample were cured, trimmed and sectioned (70-90 nm) with a Leica UC6 ultramicrotome (Leica Microsystems, Germany); grids were post-stained with 2% aqueous uranyl acetate and Reynold’s lead citrate. Transmission electron microscope imaging was conducted using either a Tecnai 12 120kV TEM (FEI, Hillsboro, OR) or a JEOL 1200EX 80kV TEM (JEOL USA, Peabody, MA); images were collected using a Gatan Ultrascan 1000 CCD camera with Digital Micrograph software (Gatan Inc., Pleasanton, CA).

### Plunge Freezing

4 ul of *Nannochloropsis* suspensions (4 x 10^7^ cells mL^−1^) were pipetted onto a 300 mesh gold Quantifoil grid with 2um holes (Quantifoil Micro Tools, GmbH) and manually blotted from the opposite side in a Vitrobot Mark IV (ThermoFisher Scientific, MA, USA) before being plunge-frozen in liquid ethane. Grids were snapped into an autoloader assembly (c-ring clipped autogrid) for stability during downstream handling steps.

### Cryo focused ion beam and scanning electron microscopy (FIB-SEM)

Frozen grids were loaded into a Linkham CMS196 cryostage (Linkham Scientific Instruments, Tadworth, England) held at −150 °C and placed onto a Zeiss LSM 880 equipped with Airyscan for fluorescence microscopy. Fluorescence and brightfield images were collected at multiple regions across two grids so that targeted cryo-lamellae could be made of cells with prominent red bodies. Following cryo-fluorescence, and detection of autofluorescence in chloroplasts and the red body, the grids were transferred under liquid nitrogen to a Leica Ace 900 (Leica Microsystems, Germany) and coated with 5 nm of platinum prior to being transferred to the Zeiss Crossbeam 540 (Zeiss, Germany). The Zeiss Crossbeam 540 was equipped with a Leica CryoStage cooled to −150 °C, used for milling and imaging of the frozen cells. Zeiss Atlas and Zen Connect software was used to navigate and map locations to be imaged for correlative cryo-fluorescence-electron microscopy. To create cryo-lamellae for cryo-electron tomography, milling was performed using a gallium ion source at an energy of 37 pA and a working distance of 5 mm. Imaging of the grid and milled lamellae was done using an Everhart-Thornley secondary electron detector at 2.0 kV. When slicing and viewing during the milling run, to generate a three-dimensional image stack, the z-depth for each mill slice was 20 nm, and the target lamella thickness was 400 nm.

### Cryo-electron tomography (cryo-ET)

FIB-milled grids were transferred to a Titan Krios electron microscope (ThermoFisher Scientific) operated at 300kV and equipped with a BIO Quantum energy filter and a K3 direct electron detector (Gatan). Full −60 to +60 bi-derectional tilt series were collected for 3 lamellae in 2 degree increments with a total dose of 100 e-/Å2 in low dose mode in SerialEM (Mastronarde, 2005). A total of 5 tilt series were aligned in IMOD (Kremer et al., 1996) using patch tracking and tomograms were reconstructed using the simultaneous iterative reconstruction technique (SIRT) algorithm. After inspection the best tomogram was selected for its quality and 2D sections as well as movies were generated in FIJI.

### Fluorescent Labeling

Calcofluor white (CFW, Fluorescent Brightener 28 / Calcofluor White M2R; Sigma F-3397 FW 960.9, dye content 90%) binds to β-glucans, and was used to fluorescently label the cellulose cell wall component (Bidhendi et al., 2020). A 500x stock was made with water, and diluted in *f* medium to the indicated final concentrations (% weight / volume). The DNA-binding dye Hoechst 33342 (AdipoGen CDX-B0030-M025, CA, USA) was used to label nuclei at a final concentration of 5 μg/mL. For the time dependent permeability experiment, the dyes stocks were introduced to the cell culture in 50 mL conical centrifuge tubes. At each time point, and for each treatment, 1.6 mL of this stained suspension was transferred to a microfuge tube and centrifuged at 2,000 rcf x 30 seconds. Cells generally adhered to the side of the tube and did not form an obvious pellet, but pipetting off 1 mL and repeating the centrifugation led to a small loose pellet. Higher centrifugation forces and times appeared to result in higher background fluorescence and abnormal cell morphology (data not shown). The remaining supernatant was removed and replaced with 0.8 mL fresh *f* medium. This resuspension was centrifuged for 2,000 rcf x 30 seconds, the supernatant removed, and the pellet resuspended in 1.6 mL fresh *f* medium.

### Bulk Fluorescence Quantification

Fluorescence measurements were collected with an Infinite M1000 Pro plate-reading spectrophotometer/fluorometer (Tecan, Switzerland). Measurements were collected in 96-well, flat optical bottom, black walled plates (Thermo Fischer Scientific 165305). From the stained and washed cell suspensions, 200 μL was pipetted into each of 7 wells as technical replicates, and fluorescence measured with the following settings: Bottom read, Mode 1 (400 Hz), 50 flashes. Chlorophyll fluorescence-ex/em 630/680 bandwidth = 10 nm each. Gain = 100. Calcofluor white- 380/475, 10 nm / 20 nm bandwidth. Gain = 50. Hoechst- 361/497. 10 nm / 10 nm bandwidth. Gain = 100. Absorbance- 750 nm. 25 flashes, 100 ms settle time. The experiment was repeated in its entirety three times. Data was compiled, plotted, and analyzed in R; simple linear regression for plotting and analysis of covariance (ANCOVA) for testing differences in slope were carried out using the averaged technical replicate values.

### Sucrose Density Gradients and Ultracentrifugation

Discontinuous sucrose gradients for further purifying shed red body particles were created by manually underlaying denser solutions of sucrose in water beneath less dense ones with thin-tipped glass Pasteur pipettes. Pre-enriched preparations of shed red bodies suspended in water were applied to the top of the gradient and centrifuged for 30 min at 40,000 rcf in an Optima XE-90 Ultracentrifuge (Beckman Coulter, USA) with SW60 Ti rotor and Beckman Ultra-Clear 11 x 60 mm centrifuge tubes (#344062).

### Generation of *CzBKTox* carotenoid mutants

*Chromochloris zofingiensis* cDNA was isolated as described in (Roth et al., 2017), and kindly provided by Daniel Westcott. From *C. zofingiensis* cDNA, *CzBKT1 (*AY772713) (Huang et al., 2006) was isolated with gene specific primers in which a 3xFLAG-tag was introduced at the 3’ end. *CzBKT1,* with and without *N. oceanica*-specific chloroplast targeting sequence (cTP) (Moog et al., 2015), was assembled with a hygromycin resistance cassette (HygR) (Vieler et al., 2012), *CAH1* promoter and *ARF* terminator (Gee and Niyogi, 2017) in pDONR221, by Gibson assembly (Invitrogen, USA). HygR-*CzBKT* constructs with and without cTP were linearized by PCR prior to transformation into *N. oceanica* (Vieler et al., 2012).

### Ultra High performance liquid chromatography (UHPLC) and HPLC analysis

UHPLC analysis was done using the method described in (Bijttebier et al., 2014). In brief, the filtered sample extract was analyzed using an Ultimate 3000 UHPLC+ Focused system (Dionex Corporation, Sunnyvale, CA, USA) coupled to a Bruker Compact APCI-QTOF-MS (Bruker) system. Samples were separated on a ACQUITY UPLC HSS C18 SB column (100 × 2.1 mm ID, 1.8 µm particle size, 100 Å pore size; WATERS; Milford, MA) maintained at 40 °C with a flow rate of 0.5 mL min-1 and mobile phase consisting of A: 50:22.5:22.5:5 water + 5 mM ammonium acetate:methanol:acetonitrile:ethyl acetate, B: 50:50 acetonitrile:ethyl acetate. For detection diode array detection (DAD) UV-Vis 190-800nm and Atmospheric pressure chemical ionization (APCI) was used as ionization technique for the mass spectroscopy (MS) detection. Briefly, the corona discharge current was set at 5 μA, the vaporizer and capillary temperature were set at 450 and 350 ◦ C, respectively and a capillary voltage of ±25 V was used for both positive and negative APCI. A scanning range from m/z 100 to m/z 1400 was used for the MS in all analytical methods. Pigment analysis by fluorescence detection by HPLC was performed as described in (Leonelli et al., 2016; Müller-Moulé et al., 2002).

### Fourier transform infrared spectroscopy (FTIR)

FTIR spectra were collected with a Vertex 80 Time-resolved FTIR (Bruker, MA, USA) in attenuated total reflectance mode (ATR). 100 spectra from 4000 to 400 cm^−1^ were collected for each sample at a resolution of 4 cm^−1^. Adhesive backed aluminum foil used for sealing 96-well PCR plates was cut into strips and adhered to a glass microscope slide for ease of handling. Aluminum provides an inexpensive substrate with low background for ATR FTIR (Cui et al., 2016). These were cleaned with ethanol and stored with desiccant until use. Concentrated “red sediment”, red body-enrichment (“red sediment” subjected to high-pressure homogenization, sucrose density gradient and water washes-see Figure 9 supplemental 1), and red body-depletion suspensions (SDS incubation of “red sediment” followed by water washes) were spotted onto the foil (5 μL) and placed in a vacuum bell with desiccant until the water was evaporated (> 30 minutes). Samples were stored in a microscope slide holder in a sealed container with desiccant at 4 °C until spectra collection later that day. A position on the foil was left empty to serve as a calibration blank prior to other measurements, and two separate sample spots for each sample type were measured with similar results.

All ATR FTIR data were subject to preprocessing that included conversion from reflectance to absorbance and baseline correction. Second derivatives of the absorption spectra were calculated as a slope through seven neighboring points using Savitsky-Golay in the OMNIC software program (version 9.8, Thermo Fisher Scientific). Peak position determination and data interpretation was guided by the second derivative spectra (Figure 9-supplemental 2), published literature, and the NIST open-source library.

## Acknowledgements

We thank Drs. Steven Ruzin and Denise Schichnes for assistance and consultations with the light microscopy presented here. We thank Joel Mancuso, Vimal Gangadharan, and Jeffrey Marshman of Zeiss Inc. and Abigail Lytton-Jean and David Mankus of the KI Nanomaterials Lab at MIT for their time, expertise, and assistance in cryo-fluorescence and cryo-FIB-milling. Valuable feedback during figure preparation was provided by many colleagues, especially Dhruv Patel, Simon Álamos, and Lorenzo Washington. We thank Dr. Daniel Westcott for providing *Chromochloris zofingiensis* cDNA, Dr. Masakazu Iwai for providing valuable training and advice regarding sucrose density gradients, Dr. Jian-Hua Chen for performing a preliminary soft x-ray tomography experiment, and Ethan Boynton for critical comments on the manuscript. This article is subject to HHMI’s Open Access to Publications policy. HHMI lab heads have previously granted a nonexclusive CC BY 4.0 license to the public and a sublicensable license to HHMI in their research articles. Pursuant to those licenses, the author-accepted manuscript of this article can be made freely available under a CC BY 4.0 license immediately upon publication.

---

**Figure 1- supplemental 1.**
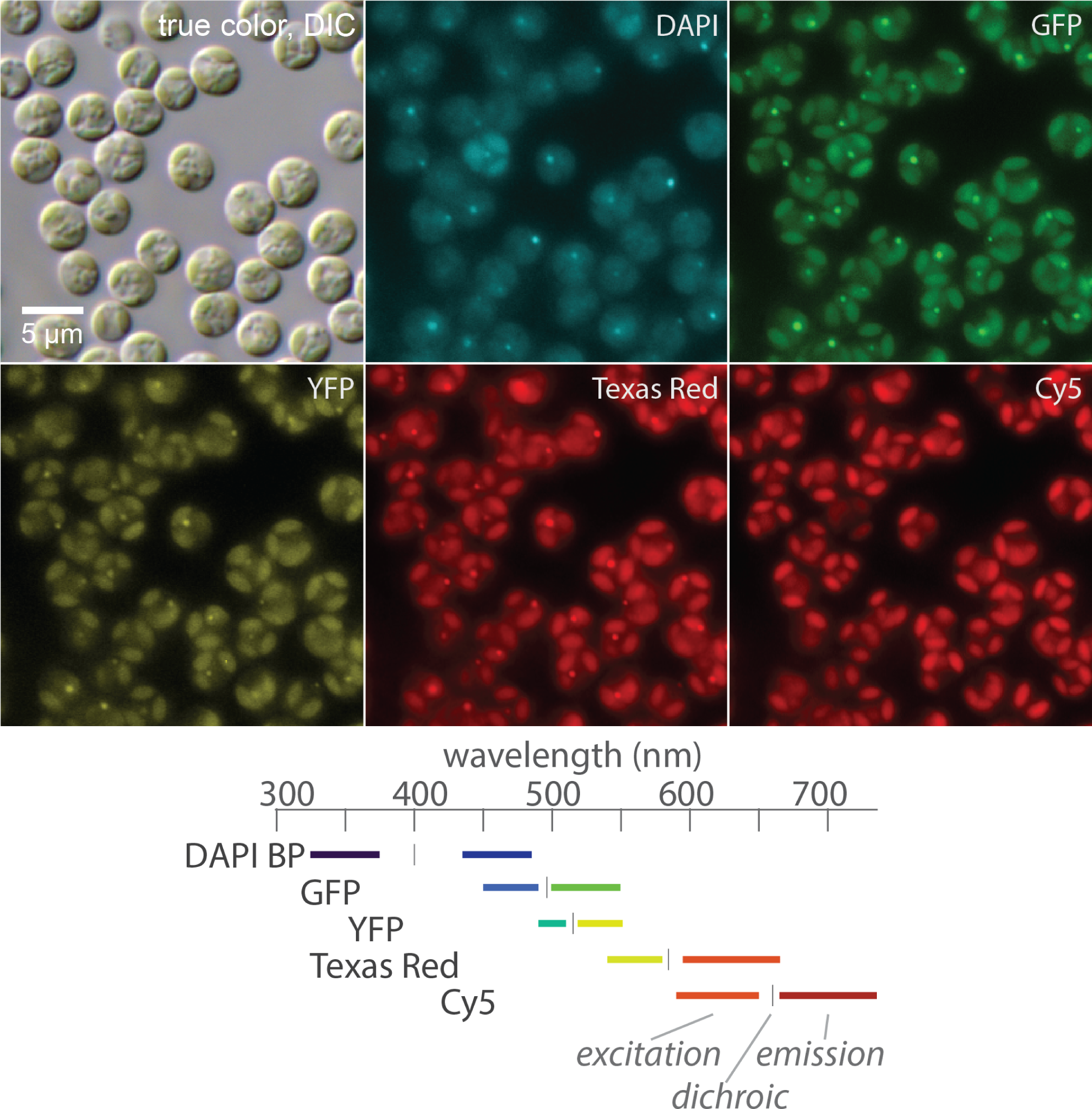
Apparent red body autofluorescence through different filter sets. Synchronously dividing cells grown in a 12-h light/12-h dark photoperiod were imaged on a wide-field fluorescence microscope. For fluorescence images, shown are 2D maximum intensity projections from three z positions that spanned the majority of the cell volumes. Channel labels refer to the filter sets used; all signal is derived from autofluorescence and false colored to approximate true color (except for DAPI, displayed in cyan for visibility). Filter set specifications are graphically depicted at the bottom.

**Figure 1- supplemental 2.**
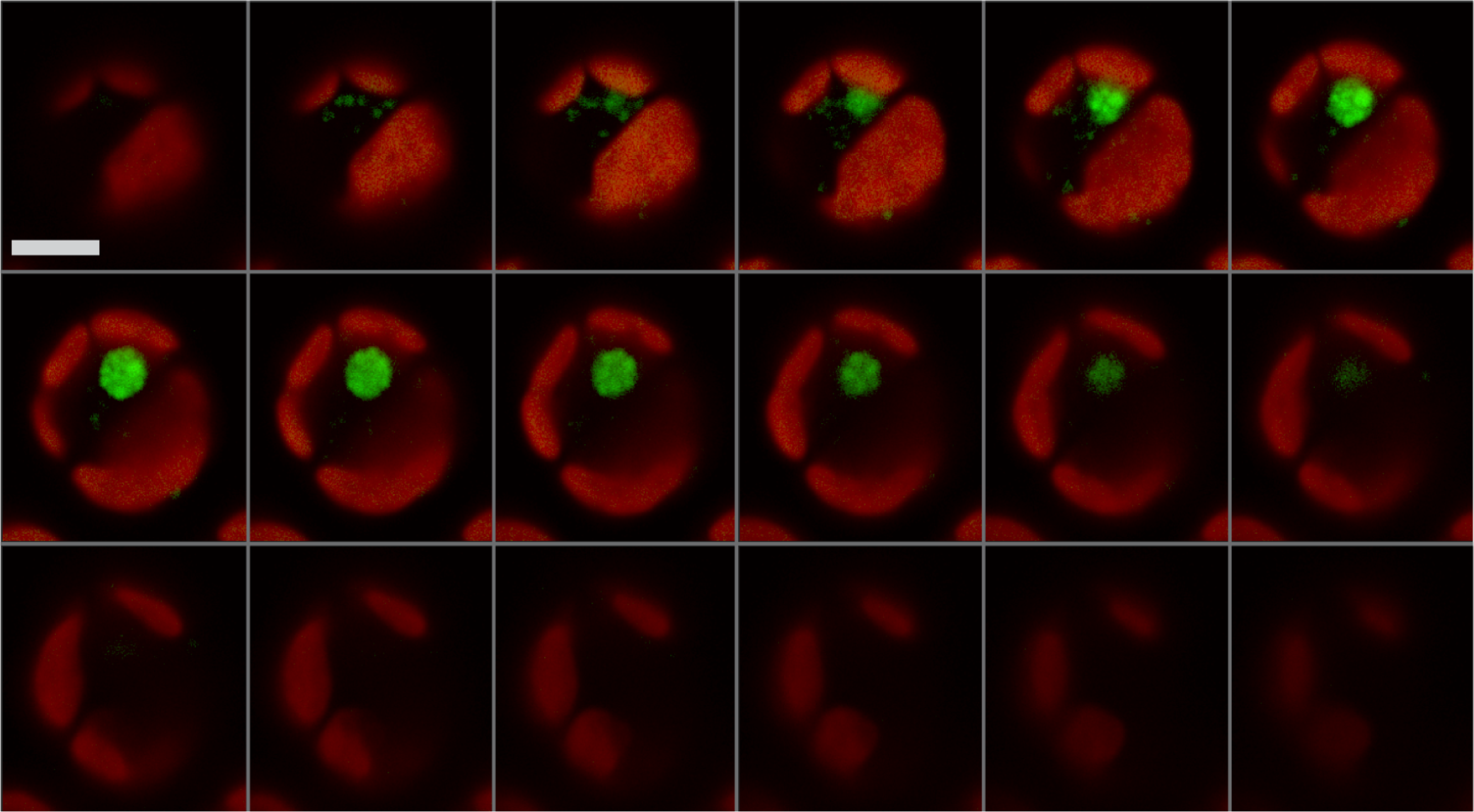
Some eustigmatophyte red bodies appear to be aggregations of smaller globules. Confocal scanning laser microscopy z-stack of a single *Vischeria* sp. cell. Green autofluorescence (pseudo colored green, ex/em: 488 nm / 525 nm), and chlorophyll autofluorescence (pseudo colored red, ex/em: 633 nm / >680 nm). Each slice shown was collected slightly less than 0.5 μm apart in the z dimension. Scale bar = 5 μm.

**Figure 2- supplemental 1.**
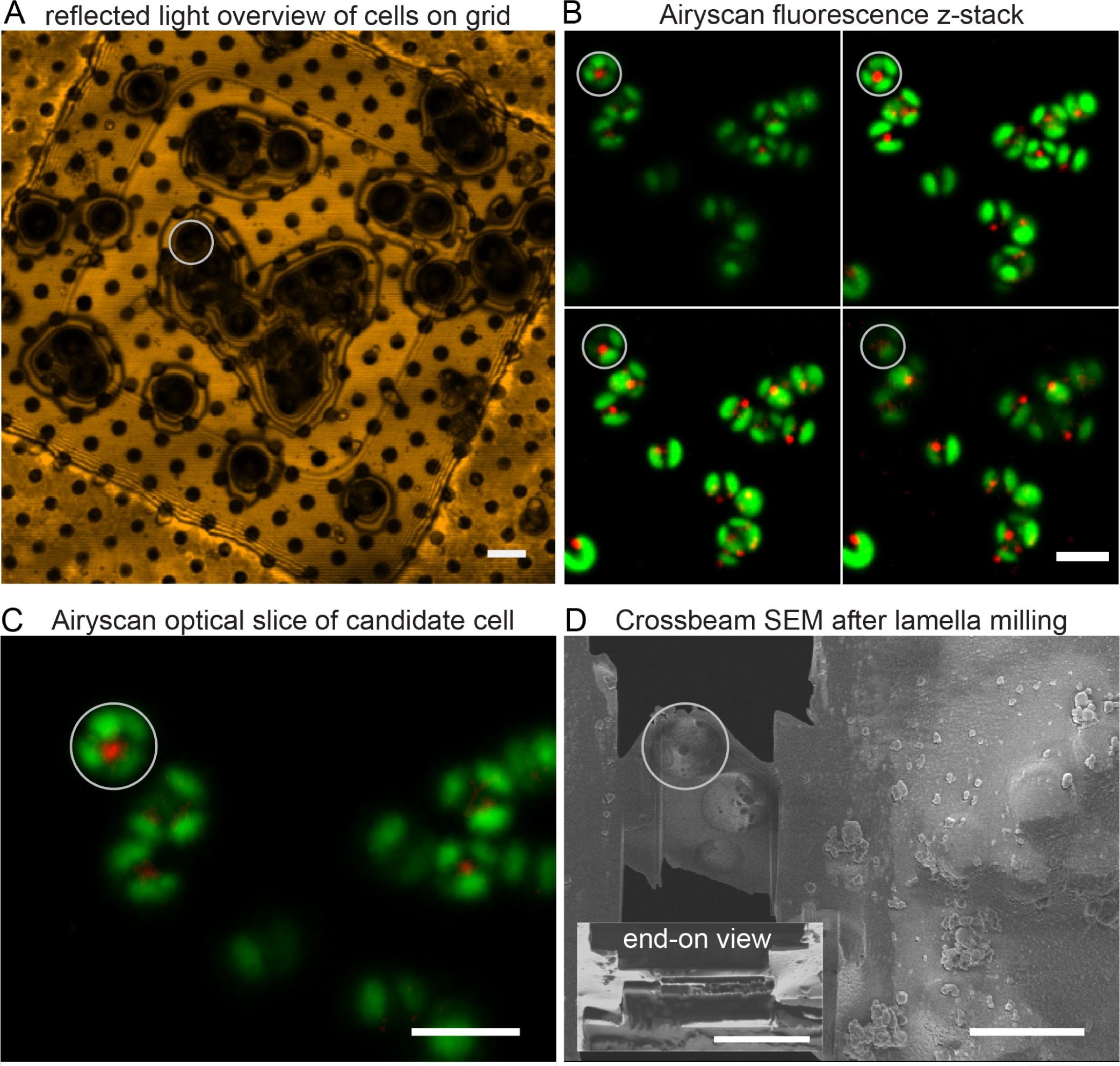
Cryo electron tomography lamellae preparation and correlative light microscopy. **(A)** A relatively low magnification “overview” image was acquired by reflected light of frozen cells on the carbon grid. **(B)** Cryo autofluorescence z-stacks were acquired to locate candidate red bodies (red false colored) among chloroplasts (green false colored) in three dimensions. A subsample of a z-stack is shown here for brevity. Note that the red/green false colors were rendered differently here than in other images in this manuscript. **(C)** Optical section of a candidate cell that contained a prominent red body (red false colored) and distinct chloroplast autofluorescence features to guide FIB-SEM, and identify features in cryo-TEM images. **(D)** SEM views of a finished physical section (lamella) after beam milling of the candidate cell in preparation for cryo TEM tomography. The main panel shows the lamella “top down”; inset shows an example lamella “end-on”. The scale bar = 5 μm in all images. The same candidate cell is circled in each image throughout the process.

**Video 1. Cryo electron tomography of a *Nannochloropsis oceanica* cell.**

Reconstruction of a 400 nm thick lamella of *Nannochloropsis oceanica*. The movie of successive tomographic sections (spaced by 4.42 nm) through the entire reconstruction shows the cellular context of a complete red body volume (spherical, electron-dense feature in center-right of frame). Scale bar = 500 nm. Presumptive assignments for organelles are annotated in the video (plastid/chloroplast, nucleus, mito/mitochondrion, CER/chloroplast endoplasmic reticulum, RB/red body), as well as in a still frame in Figure 2.

**Figure 4- supplemental 1.**
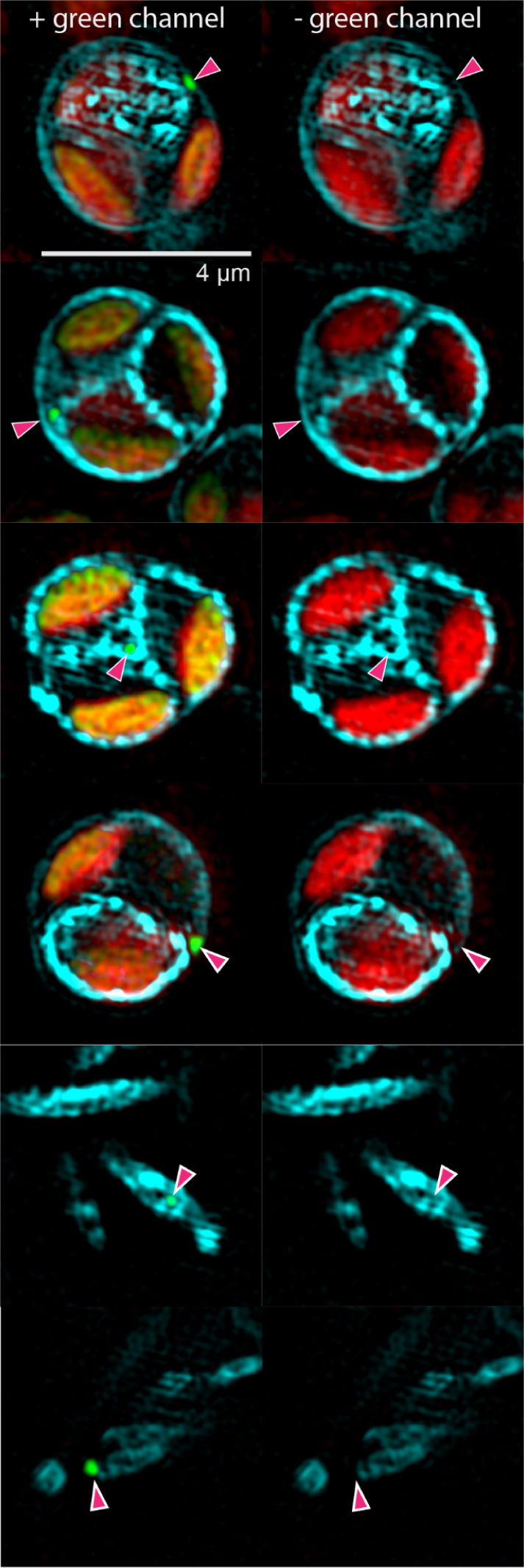
The red body resides in the apoplast during autospore cell wall formation. Synchronously dividing cells were sampled 3 hours after subjective dusk and bound to polylysine-coated coverslips. These were incubated for 1 hour in the dark with media + 0.05% calcofluor white, a cellulose-binding fluorescent dye. These cells were imaged with SIM (cyan = excitation 405 nm, emission 420-480 nm; green = excitation 488 nm, emission 495-550 nm; red = excitation 642 nm, emission >655 nm). Displayed are example cells that exhibited visibly stained cell walls, each with and without the green channel to allow examination of the area around the green fluorescence of the red body (indicated by magenta arrowhead). In some cases, what appeared to be shed autosporangial walls were visible with fluorescent punctae visible within (lower two image pairs).

**Figure 5- supplemental 1.**
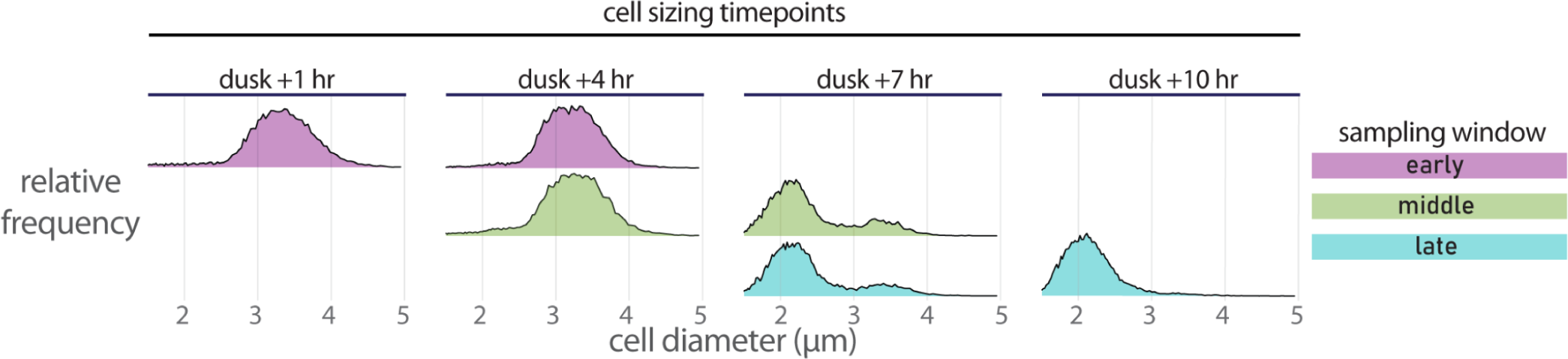
Cell sizing at beginning and end of staining windows confirms synchronous autospore release. Cell diameter was quantified with a Coulter counter at the indicated time points. Each window was sampled at the beginning and end, so the dusk +4 h and dusk +7 h time points show cell size distributions for two samplings, which can detect perturbations in division due to culture handling. Distributions are plotted as relative frequency with the same vertical scaling (bin size ∼ 0.01 μm, maximum vertical height ∼3.6%, for each distribution, n > 10k). Results are shown for one experiment out of the three performed.

**Figure 5- supplemental 2.**
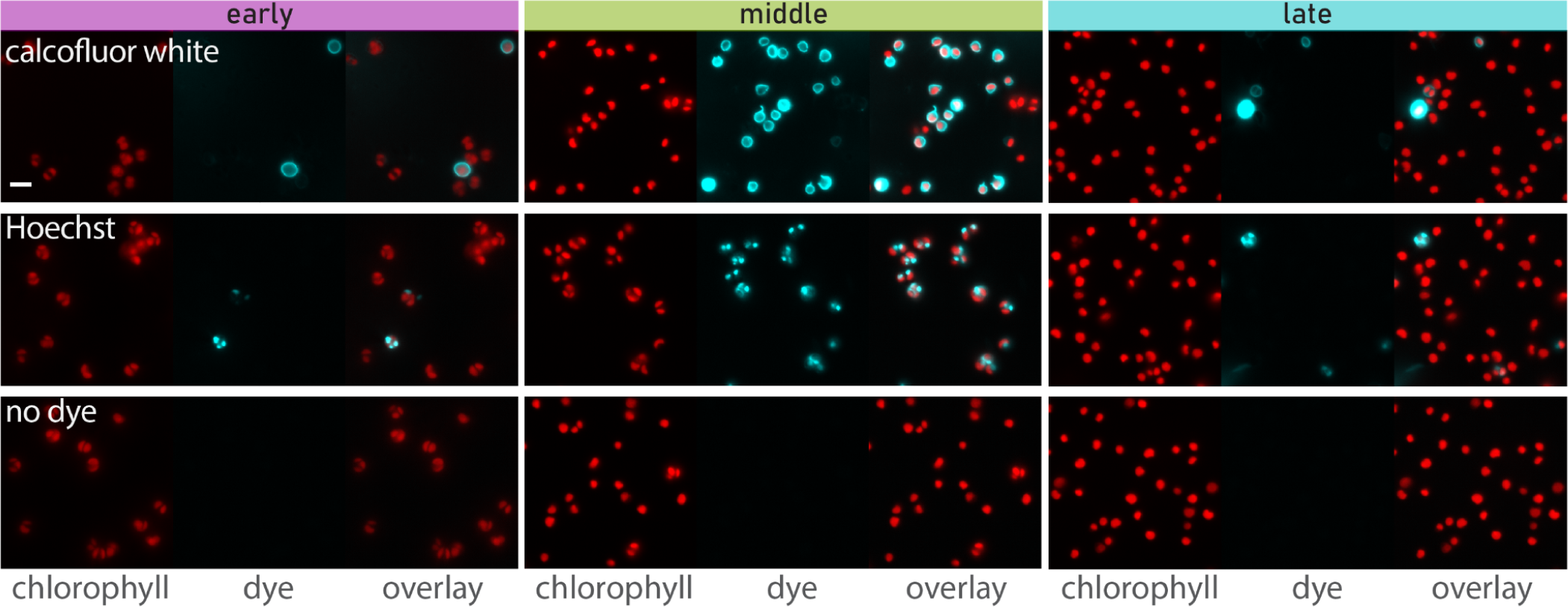
Additional microscopy of CFW and Hoechst stained cells. Images were acquired and are presented in a similar way as in Figure 5. Images are of cells from a different independent experiment from those presented in Figure 5.

**Figure 5- source data 1. Bulk Fluorescence Quantification.** This comma separated value (.csv) file contains the raw fluorescence quantitation along with measurement metadata and the normalized values. Date = date of experiment; Window = “early”, “middle”, and “late” staining periods; Minutes_Elapsed = timestamp of acquisition converted to minutes after subjective dusk; Treatment = the dye used, if applicable; Technical_Replicate = individual wells, Abs750 = absorbance at 750 nm, not used in Figure 5; Chlorophyll = chlorophyll autofluorescence (arbitrary units); Hoechst = Hoechst 33342 fluorescence (arbitrary units); Calcofluor_White = calcofluor white (CFW) fluorescence (arbitrary units); the remaining columns contain dye fluorescence divided by the Abs750 or Chlorophyll value, e.g. “Hoechst_N_Chl” is Hoechst / Chlorophyll.

**Figure 7- supplemental 1.**
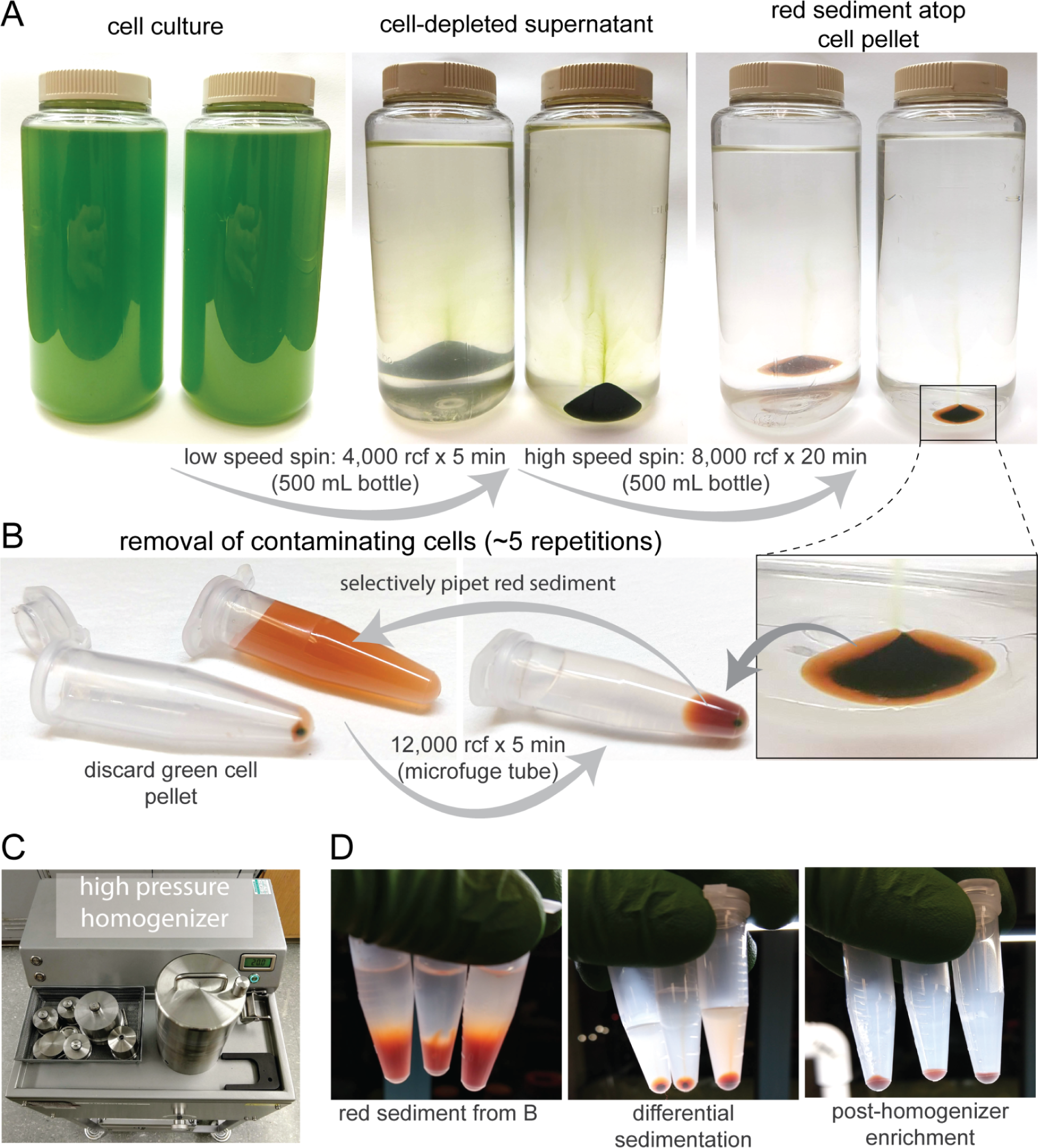
Isolation procedure of the red sediment and isolation of shed red body particles. **(A)** A procedure to isolate shed walls and red bodies from high density liter scale liquid cultures grown at 3% CO_2_ and 100 μmol photons m^−2^ s^−1^. A relatively low intensity centrifugation first depletes the supernatant of whole cells, and this supernatant is then subjected to a higher intensity centrifugation to pellet remaining cells and red sediment. **(B)** The pellet from (A) is further depleted of cells by repeated centrifugation in microfuge tubes and selective pipetting to avoid transferring the lower cell pellet to the next repetition. **(C)** The isolated red sediment from (B) is passed through a high-pressure homogenizer and recollected in microfuge tubes. **(D)** Further differential sedimentation to isolate the shed red body particles. Left panel-original red sediment as in (B), middle panel-samples after homogenizer treatment and the first round of differential sedimentation as in (B), except the dark lower pellet is retained for subsequent repetitions. Right panel-the dark pellet after several rounds of depletion of residual shed wall particles.

**Figure 7- supplemental 2.**
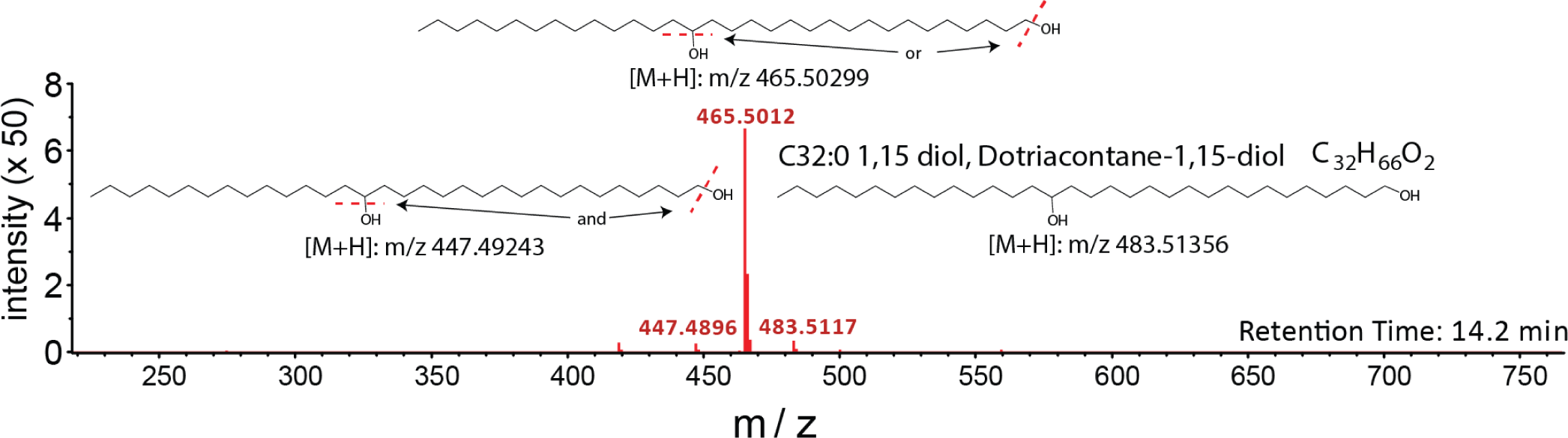
APCI In-source fragmentation of putative C_32_ long-chain diol. C_32_ long-chain diol is depicted as 1,15-dotriacontanediol based on long chain diols characterized in Balzano et al. (2019). m/z values measured are within 5 ppm of predicted m/z for the identified ions.

**Figure 8- supplemental 1.**
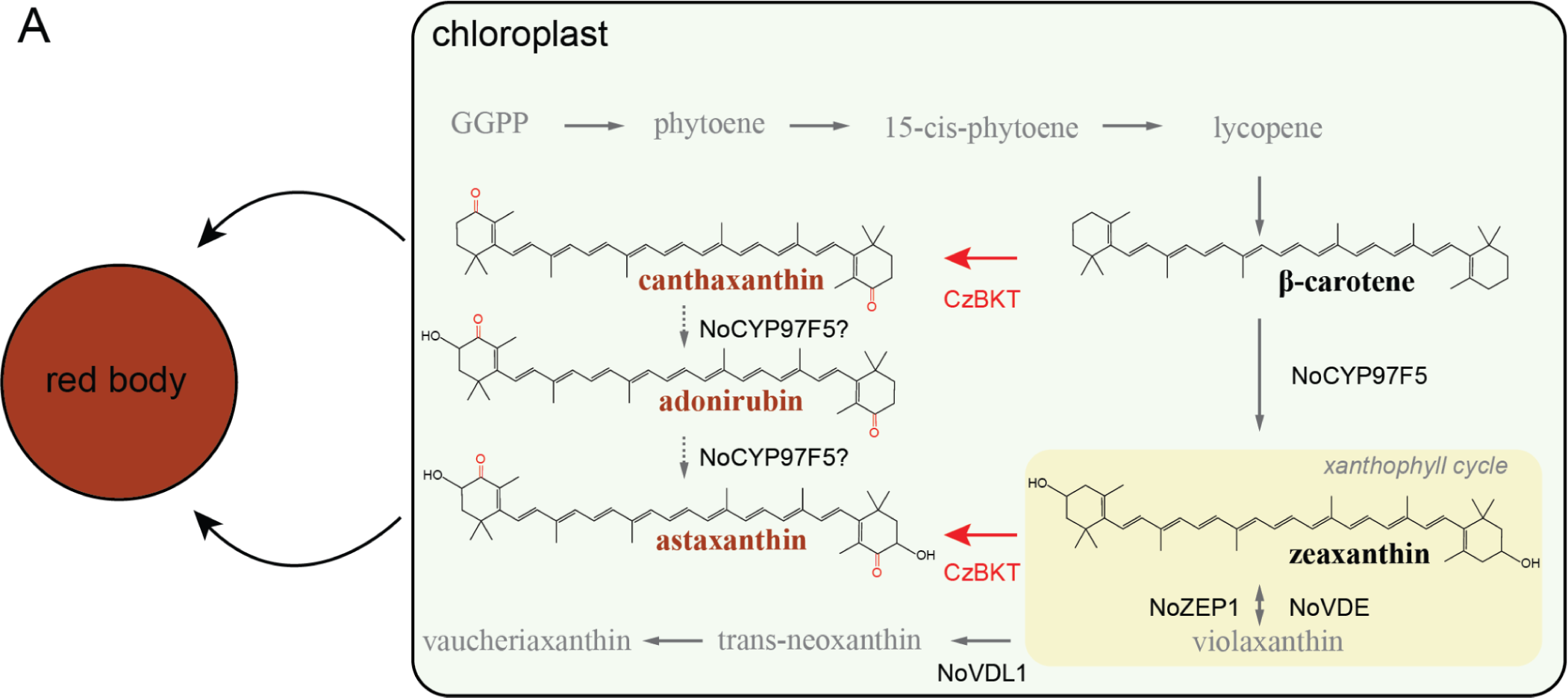
Rationale for *CzBKT* overexpression. A schematic biosynthetic pathway for carotenoids in *Nannochloropsis* is shown. While the endogenous pathway leading to canthaxanthin and astaxanthin are not precisely known (dotted lines). NoCYP97F5 have previously been shown to catalyze C3 & C3’ hydroxylation of beta-carotene (Leonelli et al., 2016), similar to the hydroxylation required for astaxanthin biosynthesis from canthaxanthin. Heterologous expression of a beta-carotene ketolase (BKT) would possibly lead to ketocarotenoid accumulation and downstream effects on the red body size and composition.

**Figure 9- supplemental 1.**
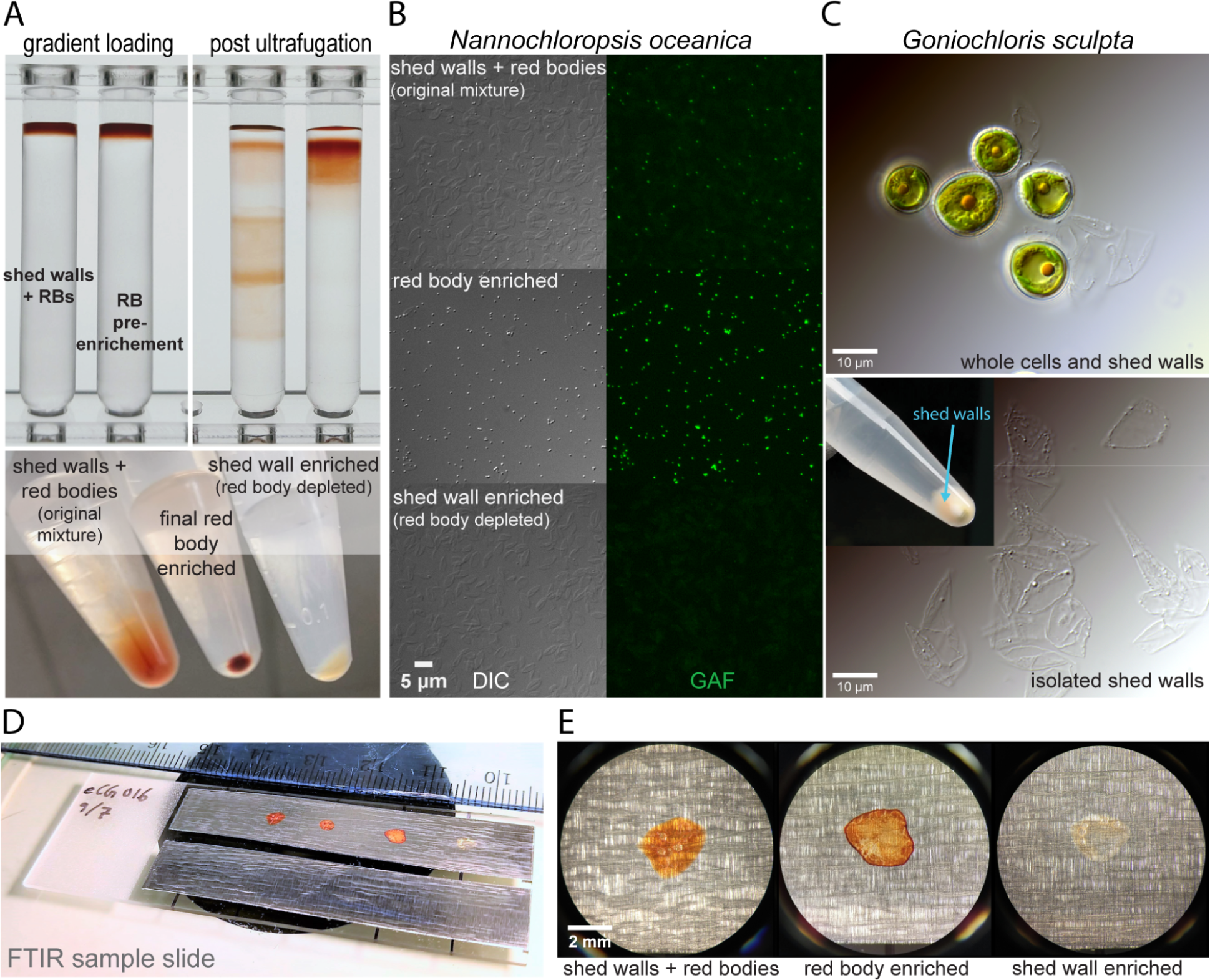
Preparation of FTIR samples. **(A)** Discontinuous sucrose density gradient to separate residual shed wall particles from the red body enrichment. Left panel-initial loading of the centrifuge tubes at the top of 20%, 26.7%, 33.3% and 40% sucrose (w/v with water). Left tube was loaded with the red sediment mixture of shed walls and red bodies, right tube with the high pressure homogenizer treated red sediment as described in Figure 7- supplemental 1. Right panel-after 30 minutes at 40,000 rcf. The final enrichment consisted of the dark band at the very top of the right tube. Bottom panel-one of the water wash steps (of 5 total) after recollection from the gradient, plus a red body depleted sample of the original mixture incubated with 1% sodium dodecyl sulfate (SDS) at 50°C for 10 min. **(B)** Quality control microscopy of the enriched fractions. Samples from (A) were bound to coverglasses and imaged as described previously. GAF = green autofluorescence. **(C)** Isolation of apparent shed walls from *G. sculpta* cultures. Upper panel-cells and shed walls from the agar culture plate. Lower panel-shed walls after differential sedimentation and selective pipetting (see inset). (**D)** Samples dried down onto aluminum foil in preparation for FTIR measurement. **(E)** View of dried samples through dissecting scope ocular. Three separate images joined to present them side by side.

**Figure 9 - supplemental 2.**
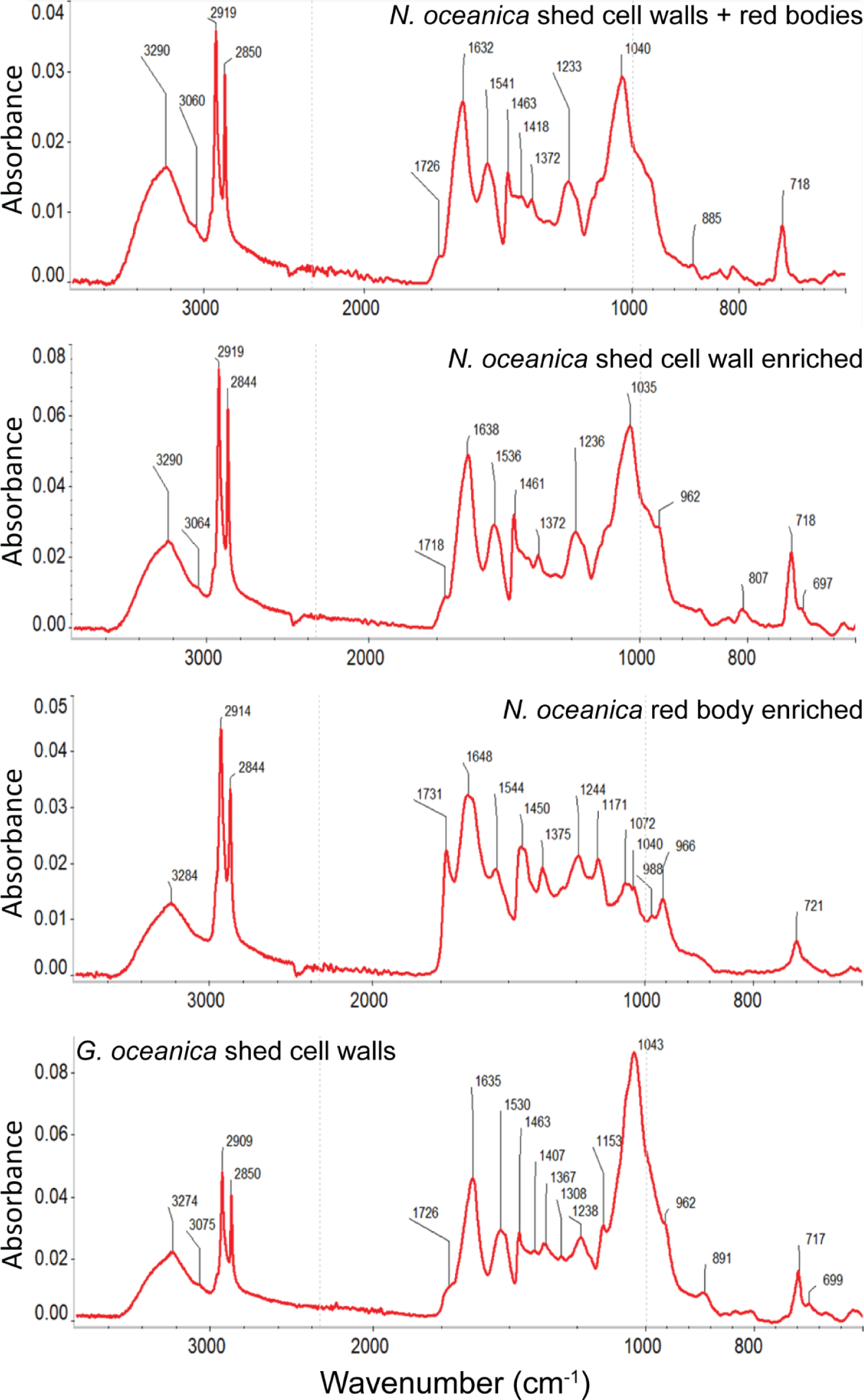
ATR-FTIR absorbance spectrum peak calls. Infrared absorbance spectra from Figure 9 are shown with local maxima annotations (in wavenumber). Additionally, included here is a spectrum for the shed cell walls of the eustigmatophyte alga, *Goniochloris sculpta*.

**Figure 9- supplemental 3.**
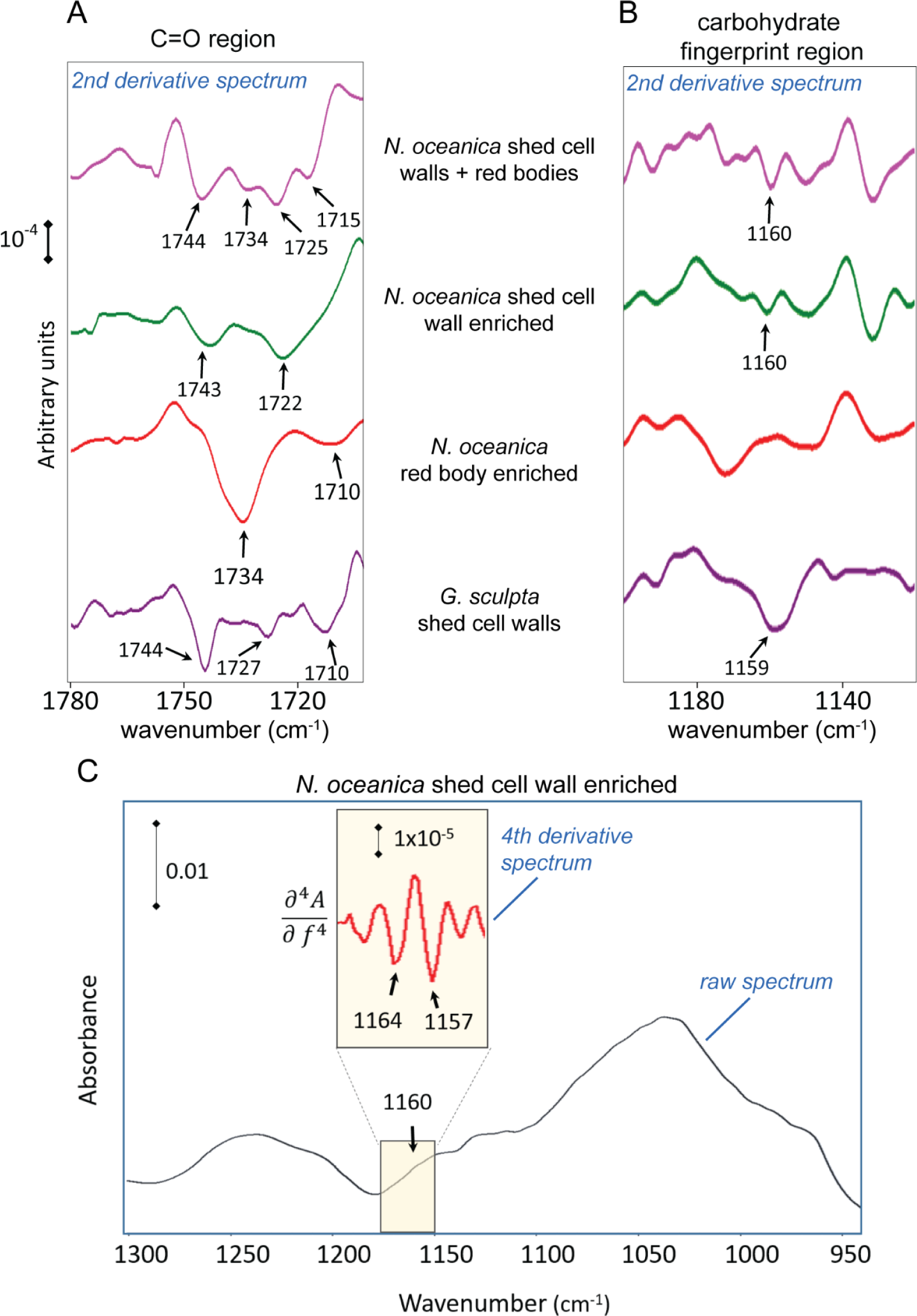
Analyses of second and higher order derivatives of ATR-FTIR spectra reveal additional information about chemical composition. The prominent bands in the raw absorption spectra if Figure 9 were identified by the corresponding second derivative spectra. **(A)** Second derivative spectra and peak calls for the non-peptide carbonyl region, and **(B)** a segment of the carbohydrate fingerprint region. **(C)** The shoulder at 1160 cm^−1^ was further decomposed by 4^th^ derivative analysis.

**Figure 9- supplemental 4.**
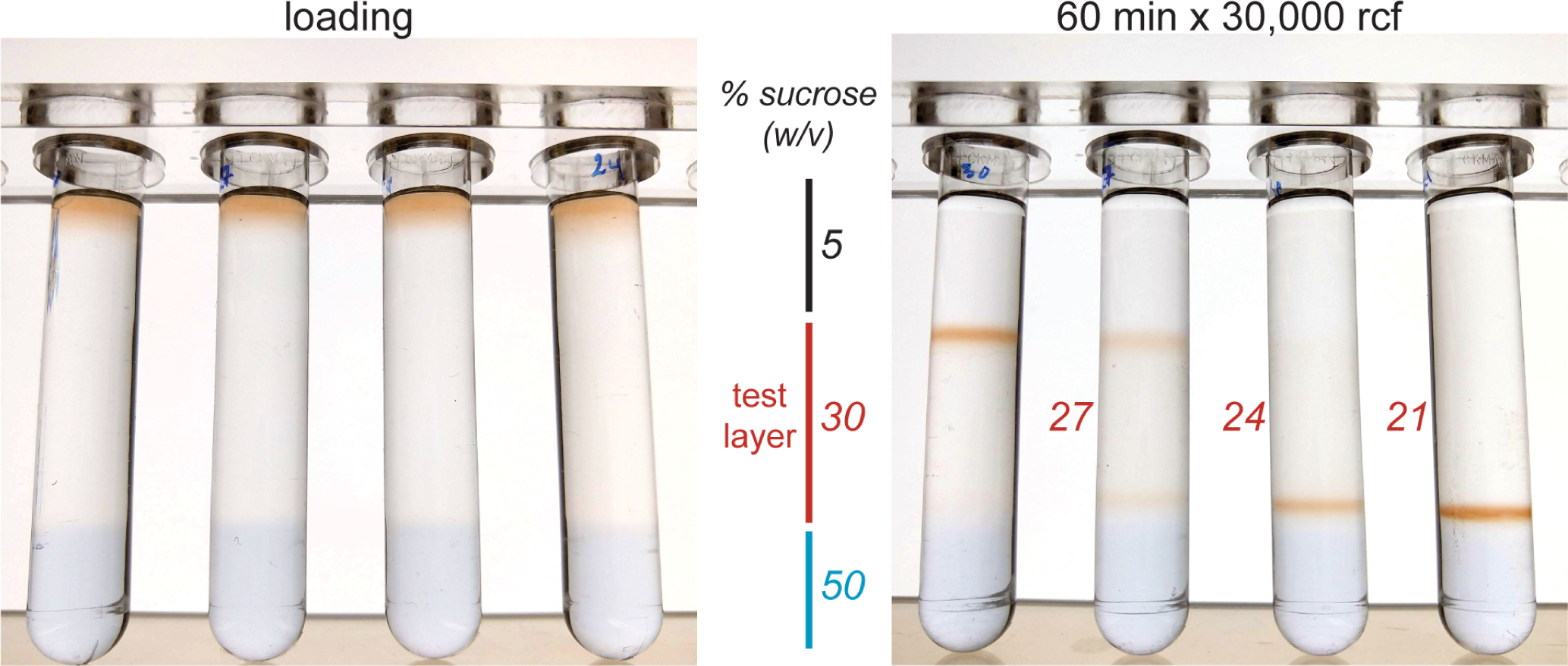
Density estimation of isolated shed red bodies by discontinuous sucrose gradients. Shed red bodies were isolated as shown in Figure 9- supplemental 1, and loaded onto sucrose step gradients composed of a 5% (w/v) focusing layer, a test layer (either 30%, 27%, 24%, or 21%), and a stopping layer (50%) made visible with a small amount of bromophenol blue. Gradients were subjected to 30,000 relative centrifugal force (rcf) for 60 min at 20°C, and examined to see if the red bodies sedimented above or below the test layer.

**Figure 9- source data 1. ATR-FTIR absorbance values.** This comma separated value (.csv) file contains the raw transmittance values for the spectra shown in Figure 9.

**Figure 10- source data 1. ANOVA and post-hoc Tukey HSD comparisons for red body volume timecourse.** This spreadsheet (.xlsx) contains the output of an analysis of variance (ANOVA) and a post-hoc Tukey honestly significant difference (HSD) test from R. Of the many comparisons possible, key comparisons showing differences within a light level over time, and differences in maximal red body volume between light levels are included in a separate tab. The Tukey HSD output includes the estimate of the difference in volume (cubic microns), the 95% confidence interval around that difference, and the p-value for that difference adjusted for multiple comparisons.

**Figure 10- source data 2. Compiled timecourse red body volumetric estimates.** This spreadsheet (.xlsx) contains the volumetric estimates from 3 dimensional reconstructions of SR-SIM image stacks, along with measurement metadata and additional quantifications like fluorescence intensity. Light = the culture growth light intensity; Time = time point in hours in relation to subjective dusk (12/12 photoperiod); Image = “a” or “b”, each Light x Time combination included two separate image stacks of different cells; Volume = volumetric estimate in cubic microns; Area = area estimation in square microns; Sphericity = the ratio of the actual area and the area of a sphere of the same volume; Intensity * = different statistics of fluorescence intensity for a given volume (arbitrary fluorescence units); Position Z = the “depth” of the object in the image stack. An additional tab includes a sample size table for each Light x Time combination. The variation in n reflects the different number of cells captured in the two image stacks.

## References

Alboresi A, Perin G, Vitulo N, Diretto G, Block M, Jouhet J, Meneghesso A, Valle G, Giuliano G, Maréchal E, Morosinotto T. 2016. Light Remodels Lipid Biosynthesis in *Nannochloropsis gaditana* by Modulating Carbon Partitioning between Organelles. Plant Physiol 171:2468–2482. doi:10.1104/pp.16.00599

Amaral R, Fawley KP, Němcová Y, Ševčíková T, Lukešová A, Fawley MW, Santos LMA, Eliáš M. 2020. Toward Modern Classification of Eustigmatophytes, Including the Description of Neomonodaceae Fam. Nov. and Three New Genera. J Phycol 56:630–648. 10.1111/jpy.12980

An G-H, Suh O-S, Kwon H-C, Kim K, Johnson EA. 2000. Quantification of carotenoids in cells of *Phaffia rhodozyma* by autofluorescence. Biotechnol Lett 22:1031–1034. doi:10.1023/A:1005614010003

Anderson NG. 1966. The Development of Zonal Centrifuges and Ancillary Systems for Tissue Fractionation and Analysis. U.S. Department of Health, Education, and Welfare, Public Health Service, National Cancer Institute.

Antia NJ, Cheng JY. 1982. The keto-carotenoids of two marine coccoid members of the Eustigmatophyceae. Br Phycol J 17:39–50. doi:10.1080/00071618200650061

Balzano S, Villanueva L, de Bar M, Sahonero Canavesi DX, Yildiz C, Engelmann JC, Marechal E, Lupette J, Sinninghe Damsté JS, Schouten S. 2019. Biosynthesis of Long Chain Alkyl Diols and Long Chain Alkenols in *Nannochloropsis spp.* (Eustigmatophyceae). Plant Cell Physiol 60:1666–1682. doi:10.1093/pcp/pcz078

Balzano S, Villanueva L, de Bar M, Sinninghe Damsté JS, Schouten S. 2017. Impact of culturing conditions on the abundance and composition of long chain alkyl diols in species of the genus *Nannochloropsis*. Org Geochem 108:9–17. doi:10.1016/j.orggeochem.2017.02.006

Barth A. 2007. Infrared spectroscopy of proteins. Biochim Biophys Acta BBA - Bioenerg 1767:1073–1101. doi:10.1016/j.bbabio.2007.06.004

Baumann WJ, Ulshöfer HW. 1968. Characteristic absorption bands and frequency shifts in the infrared spectra of naturally-occurring long-chain ethers, esters and ether esters of glycerol and various diols. Chem Phys Lipids 2:114–128. doi:10.1016/0009-3084(68)90037-6

Beisson F, Li-Beisson Y, Pollard M. 2012. Solving the puzzles of cutin and suberin polymer biosynthesis. Curr Opin Plant Biol 15:329–337. doi:10.1016/j.pbi.2012.03.003

Benito V, Goñi-de-Cerio F, Brettes P. 2015. BODIPY vital staining as a tool for flow cytometric monitoring of intracellular lipid accumulation in *Nannochloropsis gaditana*. J Appl Phycol 27:233–241. doi:10.1007/s10811-014-0310-x

Bijttebier S, D’Hondt E, Noten B, Hermans N, Apers S, Voorspoels S. 2014. Ultra high performance liquid chromatography versus high performance liquid chromatography: Stationary phase selectivity for generic carotenoid screening. J Chromatogr A 1332:46–56. doi:10.1016/j.chroma.2014.01.042

Bína D, Gardian Z, Herbstová M, Litvín R. 2017. Modular antenna of photosystem I in secondary plastids of red algal origin: a *Nannochloropsis oceanica* case study. Photosynth Res 131:255–266. doi:10.1007/s11120-016-0315-1

Bišová K, Zachleder V. 2014. Cell-cycle regulation in green algae dividing by multiple fission. J Exp Bot 65:2585–2602. doi:10.1093/jxb/ert466

Boeynaems S, Alberti S, Fawzi NL, Mittag T, Polymenidou M, Rousseau F, Schymkowitz J, Shorter J, Wolozin B, Van Den Bosch L, Tompa P, Fuxreiter M. 2018. Protein Phase Separation: A New Phase in Cell Biology. Trends Cell Biol 28:420–435. doi:10.1016/j.tcb.2018.02.004

Britton G, Liaaen-Jensen S, Pfander H. 2004. Carotenoids, 1st ed. Birkhäuser Basel.

Burczyk J, Szkawran H, Zontek I, Czygan F-Ch. 1981. Carotenoids in the outer cell-wall layer of *Scenedesmus* (Chlorophyceae). Planta 151:247–250. doi:10.1007/BF00395176

Cano M, Karns DAJ, Weissman JC, Heinnickel ML, Posewitz MC. 2021. Pigment modulation in response to irradiance intensity in the fast-growing alga *Picochlorum celeri*. Algal Res 58:102370. doi:10.1016/j.algal.2021.102370

Carbonera D, Agostini A, Di Valentin M, Gerotto C, Basso S, Giacometti GM, Morosinotto T. 2014. Photoprotective sites in the violaxanthin–chlorophyll a binding Protein (VCP) from *Nannochloropsis gaditana*. Biochim Biophys Acta BBA - Bioenerg 1837:1235–1246. doi:10.1016/j.bbabio.2014.03.014

Chaturvedi R, Fujita Y. 2006. Isolation of enhanced eicosapentaenoic acid producing mutants of *Nannochloropsis oculata* ST-6 using ethyl methane sulfonate induced mutagenesis techniques and their characterization at mRNA transcript level. Phycol Res 54:208–219. doi:10.1111/j.1440-1835.2006.00428.x

Corteggiani Carpinelli E, Telatin A, Vitulo N, Forcato C, D’Angelo M, Schiavon R, Vezzi A, Giacometti GM, Morosinotto T, Valle G. 2014. Chromosome Scale Genome Assembly and Transcriptome Profiling of *Nannochloropsis gaditana* in Nitrogen Depletion. Mol Plant 7:323–335. doi:10.1093/mp/sst120

Cui L, Butler HJ, Martin-Hirsch PL, Martin FL. 2016. Aluminium foil as a potential substrate for ATR-FTIR, transflection FTIR or Raman spectrochemical analysis of biological specimens. Anal Methods 8:481–487. doi:10.1039/C5AY02638E

Davis RW, Jones HDT, Collins AM, Ricken JB, Sinclair MB, Timlin JA, Singh S. 2014. Label-free measurement of algal triacylglyceride production using fluorescence hyperspectral imaging. Algal Res 5:181–189. doi:10.1016/j.algal.2013.11.010

Derenne S, Largeau C. 2001. A review of some important families of refractory macromolecules: composition, origin, and fate in soils and sediments. Soil Sci 166:833–847.

Ding Y, Zhang S, Yang L, Na H, Zhang P, Zhang H, Wang Y, Chen Y, Yu J, Huo C, Xu S, Garaiova M, Cong Y, Liu P. 2013. Isolating lipid droplets from multiple species. Nat Protoc 8:43–51. doi:10.1038/nprot.2012.142

Doan T-TY, Obbard JP. 2011. Improved Nile Red staining of *Nannochloropsis sp*. J Appl Phycol 23:895–901. doi:10.1007/s10811-010-9608-5

Dolch L-J, Rak C, Perin G, Tourcier G, Broughton R, Leterrier M, Morosinotto T, Tellier F, Faure J-D, Falconet D, Jouhet J, Sayanova O, Beaudoin F, Maréchal E. 2017. A Palmitic Acid Elongase Affects Eicosapentaenoic Acid and Plastidial Monogalactosyldiacylglycerol Levels in *Nannochloropsis*. Plant Physiol 173:742–759. doi:10.1104/pp.16.01420

Domínguez E, Heredia-Guerrero JA, Heredia A. 2015. Plant cutin genesis: unanswered questions. Trends Plant Sci 20:551–558. doi:10.1016/j.tplants.2015.05.009

Dunker S, Wilhelm C. 2018. Cell Wall Structure of Coccoid Green Algae as an Important Trade-Off Between Biotic Interference Mechanisms and Multidimensional Cell Growth. Front Microbiol 9:719. doi:10.3389/fmicb.2018.00719

Eliáš M, Amaral R, Fawley KP, Fawley MW, Němcová Y, Neustupa J, Přibyl P, Santos LMA, Ševčíková T. 2017. Eustigmatophyceae In: Archibald JM, Simpson AGB, Slamovits CH, editors. Handbook of the Protists. Springer International Publishing. pp. 367–406. doi:10.1007/978-3-319-28149-0_39

Fawley KP, Eliáš M, Fawley MW. 2014. The diversity and phylogeny of the commercially important algal class Eustigmatophyceae, including the new clade Goniochloridales. J Appl Phycol 26:1773–1782. doi:10.1007/s10811-013-0216-z

Fawley KP, Fawley MW. 2007. Observations on the Diversity and Ecology of Freshwater *Nannochloropsis* (Eustigmatophyceae), with Descriptions of New Taxa. Protist 158:325–336. doi:10.1016/j.protis.2007.03.003

Fawley MW, Fawley KP. 2017. Rediscovery of *Tetraedriella subglobosa* Pascher, a member of the Eustigmatophyceae. Fottea 17:96–102. doi:10.5507/fot.2016.018

Fawley MW, Jameson I, Fawley KP. 2015. The phylogeny of the genus *Nannochloropsis* (Monodopsidaceae, Eustigmatophyceae), with descriptions of *N. australis sp. nov*. and *Microchloropsis gen. nov*. Phycologia 54:545–552. doi:10.2216/15-60.1

Flori S, Jouneau P-H, Bailleul B, Gallet B, Estrozi LF, Moriscot C, Bastien O, Eicke S, Schober A, Bártulos CR, Maréchal E, Kroth PG, Petroutsos D, Zeeman S, Breyton C, Schoehn G, Falconet D, Finazzi G. 2017. Plastid thylakoid architecture optimizes photosynthesis in diatoms. Nat Commun 8:15885. doi:10.1038/ncomms15885

Gao B, Huang L, Wang F, Zhang C. 2019. *Trachydiscus guangdongensis* sp. nov., a new member of Eustigmatophyceae (Stramenopiles) isolated from China: morphology, phylogeny, fatty acid profile, pigment, and cell wall composition. Hydrobiologia 835:37–47. doi:10.1007/s10750-019-3925-8

Gao B, Yang J, Lei X, Xia S, Li A, Zhang C. 2016. Characterization of cell structural change, growth, lipid accumulation, and pigment profile of a novel oleaginous microalga, Vischeria stellata (Eustigmatophyceae), cultured with different initial nitrate supplies. J Appl Phycol 28:821–830. doi:10.1007/s10811-015-0626-1

Gee CW, Niyogi KK. 2017. The carbonic anhydrase CAH1 is an essential component of the carbon-concentrating mechanism in *Nannochloropsis oceanica*. Proc Natl Acad Sci 114:4537–4542. doi:10.1073/pnas.1700139114

Gelin F, Boogers I, Noordeloos AAM, Damsté JSS, Hatcher PG, Leeuw JW de. 1996. Novel, resistant microalgal polyethers: An important sink of organic carbon in the marine environment? Geochim Cosmochim Acta 60:1275–1280. doi:10.1016/0016-7037(96)00038-5

Gelin F, Volkman JK, De Leeuw JW, Sinninghe Damsté JS. 1997. Mid-chain hydroxy long-chain fatty acids in microalgae from the genus *Nannochloropsis*. Phytochemistry 45:641–646. doi:10.1016/S0031-9422(97)00068-X

Gelin F, Volkman JK, Largeau C, Derenne S, Sinninghe Damsté JS, De Leeuw JW. 1999. Distribution of aliphatic, nonhydrolyzable biopolymers in marine microalgae. Org Geochem 30:147–159. doi:10.1016/S0146-6380(98)00206-X

Gibbs SP. 1979. The route of entry of cytoplasmically synthesized proteins into chloroplasts of algae possessing chloroplast ER. J Cell Sci 35:253–266.

Goodenough U, Lin H, Lee J-H. 2007. Sex determination in *Chlamydomonas*. Semin Cell Dev Biol, Signal Transduction in the Myometrium 18:350–361. doi:10.1016/j.semcdb.2007.02.006

Guillard RRL, Ryther JH. 1962. Studies of Marine Planktonic Diatoms. Can J Microbiol 8:229–239. doi:10.1139/m62-029

Heimerl N, Hommel E, Westermann M, Meichsner D, Lohr M, Hertweck C, Grossman AR, Mittag M, Sasso S. 2018. A giant type I polyketide synthase participates in zygospore maturation in *Chlamydomonas reinhardtii*. Plant J 95:268–281. 10.1111/tpj.13948

Hibberd DJ. 1981. Notes on the taxonomy and nomenclature of the algal classes Eustigmatophyceae and Tribophyceae (synonym Xanthophyceae). Bot J Linn Soc 82:93–119. 10.1111/j.1095-8339.1981.tb00954.x

Hibberd DJ, Leedale GF. 1970. Eustigmatophyceae—a New Algal Class with Unique Organization of the Motile Cell. Nature 225:758–760. doi:10.1038/225758b0

Higgins HG, Stewart CM, Harrington KJ. 1961. Infrared spectra of cellulose and related polysaccharides. J Polym Sci 51:59–84. doi:10.1002/pol.1961.1205115505

Huang J-C, Wang Y, Sandmann G, Chen F. 2006. Isolation and characterization of a carotenoid oxygenase gene from *Chlorella zofingiensis* (Chlorophyta). Appl Microbiol Biotechnol 71:473–479. doi:10.1007/s00253-005-0166-8

Huertas IE, Espie GS, Colman B, Lubian LM. 2000. Light-dependent bicarbonate uptake and CO_2_ efflux in the marine microalga *Nannochloropsis gaditana*. Planta 211:43–49. doi:10.1007/s004250000254

Hyka P, Lickova S, Přibyl P, Melzoch K, Kovar K. 2013. Flow cytometry for the development of biotechnological processes with microalgae. Biotechnol Adv, Prague Symposium 2011 31:2–16. doi:10.1016/j.biotechadv.2012.04.007

Jinkerson RE, Radakovits R, Posewitz MC. 2013. Genomic insights from the oleaginous model alga *Nannochloropsis gaditana*. Bioengineered 4:37–43. doi:10.4161/bioe.21880

Jo MJ, Hur SB. 2015. Growth and Nutritional Composition of Eustigmatophyceae *Monodus subterraneus* and *Nannochloropsis oceanica* in Autotrophic and Mixotrophic Culture. Ocean Polar Res 37:61–71. doi:10.4217/OPR.2015.37.1.061

Jones RN. 1962. The effects of chain length on the infrared spectra of fatty acids and methyl esters. Can J Chem 40:321–333. doi:10.1139/v62-050

Keşan G, Litvín R, Bína D, Durchan M, Šlouf V, Polívka T. 2016. Efficient light-harvesting using non-carbonyl carotenoids: Energy transfer dynamics in the VCP complex from *Nannochloropsis oceanica*. Biochim Biophys Acta BBA - Bioenerg 1857:370–379. doi:10.1016/j.bbabio.2015.12.011

Kilian O, Benemann CSE, Niyogi KK, Vick B. 2011. High-efficiency homologous recombination in the oil-producing alga *Nannochloropsis* sp. Proc Natl Acad Sci 108:21265–21269. doi:10.1073/pnas.1105861108

Kim SS, Douglas CJ. 2013. Sporopollenin monomer biosynthesis in *Arabidopsis*. J Plant Biol 56:1–6. doi:10.1007/s12374-012-0385-3

Kleinegris DMM, van Es MA, Janssen M, Brandenburg WA, Wijffels RH. 2010. Carotenoid fluorescence in *Dunaliella salina*. J Appl Phycol 22:645–649. doi:10.1007/s10811-010-9505-y

Kremer JR, Mastronarde DN, McIntosh JR. 1996. Computer Visualization of Three-Dimensional Image Data Using IMOD. J Struct Biol 116:71–76. doi:10.1006/jsbi.1996.0013

Largeau C, Derenne S, Casadevall E, Berkaloff C, Corolleur M, Lugardon B, Raynaud JF, Connan J. 1990. Occurrence and origin of “ultralaminar” structures in “amorphous” kerogens of various source rocks and oil shales. Org Geochem 16:889–895. doi:10.1016/0146-6380(90)90125-J

Larkin P. 2011. Illustrated IR and Raman Spectra Demonstrating Important Functional Groups. Infrared and Raman Spectroscopy. Elsevier. pp. 135–176. doi:10.1016/B978-0-12-386984-5.10008-4

Leonelli L, Erickson E, Lyska D, Niyogi KK. 2016. Transient expression in *Nicotiana benthamiana* for rapid functional analysis of genes involved in non-photochemical quenching and carotenoid biosynthesis. Plant J 88:375–386. doi:10.1111/tpj.13268

Lewis RNAH, McElhaney RN. 2013. Membrane lipid phase transitions and phase organization studied by Fourier transform infrared spectroscopy. Biochim Biophys Acta BBA - Biomembr, FTIR in membrane proteins and peptide studies 1828:2347–2358. doi:10.1016/j.bbamem.2012.10.018

Ma Y, Wang Z, Yu C, Yin Y, Zhou G. 2014. Evaluation of the potential of 9 *Nannochloropsis* strains for biodiesel production. Bioresour Technol 167:503–509. doi:10.1016/j.biortech.2014.06.047

Mastronarde DN. 2005. Automated electron microscope tomography using robust prediction of specimen movements. J Struct Biol 152:36–51. doi:10.1016/j.jsb.2005.07.007

McDonald K, Müller-Reichert T. 2002. Cryomethods for thin section electron microscopy. Methods in Enzymology, Guide to Yeast Genetics and Molecular and Cell Biology Part C. Academic Press. pp. 96–123. doi:10.1016/S0076-6879(02)51843-7

McDONALD KL, Webb RI. 2011. Freeze substitution in 3 hours or less. J Microsc 243:227–233. 10.1111/j.1365-2818.2011.03526.x

Moog D, Stork S, Reislöhner S, Grosche C, Maier U-G. 2015. In vivo Localization Studies in the Stramenopile Alga *Nannochloropsis oceanica*. Protist 166:161–171. doi:10.1016/j.protis.2015.01.003

Müller-Moulé P, Conklin PL, Niyogi KK. 2002. Ascorbate deficiency can limit violaxanthin de-epoxidase activity in vivo. Plant Physiol 128:970–977. doi:10.1104/pp.010924

Murakami R, Hashimoto H. 2009. Unusual Nuclear Division in *Nannochloropsis oculata* (Eustigmatophyceae, Heterokonta) which May Ensure Faithful Transmission of Secondary Plastids. Protist 160:41–49. doi:10.1016/j.protis.2008.09.002

Nakayama T, Nakamura A, Yokoyama A, Shiratori T, Inouye I, Ishida K. 2015. Taxonomic study of a new eustigmatophycean alga, *Vacuoliviride crystalliferum* gen. et sp. nov. J Plant Res 128:249–257. doi:10.1007/s10265-014-0686-3

Ota S, Morita A, Ohnuki S, Hirata A, Sekida S, Okuda K, Ohya Y, Kawano S. 2018. Carotenoid dynamics and lipid droplet containing astaxanthin in response to light in the green alga *Haematococcus pluvialis*. Sci Rep 8:5617. doi:10.1038/s41598-018-23854-w

Park S, Steen CJ, Lyska D, Fischer AL, Endelman B, Iwai M, Niyogi KK, Fleming GR. 2019. Chlorophyll–carotenoid excitation energy transfer and charge transfer in *Nannochloropsis oceanica* for the regulation of photosynthesis. Proc Natl Acad Sci 116:3385–3390. doi:10.1073/pnas.1819011116

Parkash S, Blanshard JMV. 1975. Infrared spectra of selected ultra-pure triglycerides. Spectrochim Acta Part Mol Spectrosc 31:951–957. doi:10.1016/0584-8539(75)80158-9

Philippe G, Sørensen I, Jiao C, Sun X, Fei Z, Domozych DS, Rose JK. 2020. Cutin and suberin: assembly and origins of specialized lipidic cell wall scaffolds. Curr Opin Plant Biol, Physiology and Metabolism 55:11–20. doi:10.1016/j.pbi.2020.01.008

Poirier V, Roumet C, Munson AD. 2018. The root of the matter: Linking root traits and soil organic matter stabilization processes. Soil Biol Biochem 120:246–259. doi:10.1016/j.soilbio.2018.02.016

Poliner E, Panchy N, Newton L, Wu G, Lapinsky A, Bullard B, Zienkiewicz A, Benning C, Shiu S-H, Farré EM. 2015. Transcriptional coordination of physiological responses in *Nannochloropsis oceanica* CCMP1779 under light/dark cycles. Plant J 83:1097–1113. doi:10.1111/tpj.12944

Poliner E, Pulman JA, Zienkiewicz K, Childs K, Benning C, Farré EM. 2018. A toolkit for *Nannochloropsis oceanica* CCMP1779 enables gene stacking and genetic engineering of the eicosapentaenoic acid pathway for enhanced long-chain polyunsaturated fatty acid production. Plant Biotechnol J 16:298–309. doi:10.1111/pbi.12772

Poliner E, Takeuchi T, Du Z-Y, Benning C, Farré EM. 2019. Non-transgenic marker-free gene disruption by an episomal CRISPR system in the oleaginous microalga, *Nannochloropsis oceanica* CCMP1779. Plant J Cell Mol Biol 99:112–127. doi:10.1111/tpj.14314

Poveda-Huertes D, Patwari P, Günther J, Fabris M, Andersen-Ranberg J. 2023. Novel transformation strategies improve efficiency up to 10-fold in stramenopile algae. Algal Res 74:103165. doi:10.1016/j.algal.2023.103165

Přibyl P, Eliáš M, Cepák V, Lukavský J, Kaštánek P. 2012. Zoosporogenesis, Morphology, Ultrastructure, Pigment Composition, and Phylogenetic Position of *Trachydiscus Minutus* (Eustigmatophyceae, Heterokontophyta). J Phycol 48:231–242. 10.1111/j.1529-8817.2011.01109.x

Quilichini TD, Grienenberger E, Douglas CJ. 2015. The biosynthesis, composition and assembly of the outer pollen wall: A tough case to crack. Phytochemistry 113:170–182. doi:10.1016/j.phytochem.2014.05.002

Rabbani S, Beyer P, Kleinig H. 1998. Induced Beta-Carotene Synthesis Driven by Triacylglycerol Deposition in the Unicellular Alga *Dunaliella bardawil*. Plant Physiol 116:1239–1248.

Radakovits R, Jinkerson RE, Fuerstenberg SI, Tae H, Settlage RE, Boore JL, Posewitz MC. 2012. Draft genome sequence and genetic transformation of the oleaginous alga *Nannochloropsis gaditana*. Nat Commun 3:686. doi:10.1038/ncomms1688

Rampen SW, Datema M, Rodrigo-Gámiz M, Schouten S, Reichart G-J, Sinninghe Damsté JS. 2014. Sources and proxy potential of long chain alkyl diols in lacustrine environments. Geochim Cosmochim Acta 144:59–71. doi:10.1016/j.gca.2014.08.033

Rampen SW, Willmott V, Kim J-H, Uliana E, Mollenhauer G, Schefuß E, Sinninghe Damsté JS, Schouten S. 2012. Long chain 1,13- and 1,15-diols as a potential proxy for palaeotemperature reconstruction. Geochim Cosmochim Acta 84:204–216. doi:10.1016/j.gca.2012.01.024

Rodolfi L, Zittelli GC, Barsanti L, Rosati G, Tredici MR. 2003. Growth medium recycling in *Nannochloropsis* sp. mass cultivation. Biomol Eng 20:243–248. doi:10.1016/S1389-0344(03)00063-7

Rodriguez MC, Noseda MD, Cerezo AS. 1999. The Fibrillar Polysaccharides and their Linkage to Algaenan in the Trilaminar Layer of the Cell Wall of *Coelastrum sphaericum* (Chlorophyceae). J Phycol 35:1025–1031. doi:10.1046/j.1529-8817.1999.3551025.x

Roth MS, Cokus SJ, Gallaher SD, Walter A, Lopez D, Erickson E, Endelman B, Westcott D, Larabell CA, Merchant SS, Pellegrini M, Niyogi KK. 2017. Chromosome-level genome assembly and transcriptome of the green alga *Chromochloris zofingiensis* illuminates astaxanthin production. Proc Natl Acad Sci 114:E4296–E4305. doi:10.1073/pnas.1619928114

Ryu AJ, Kang NK, Jeon S, Hur DH, Lee EM, Lee DY, Jeong B, Chang YK, Jeong KJ. 2020. Development and characterization of a *Nannochloropsis* mutant with simultaneously enhanced growth and lipid production. Biotechnol Biofuels 13. doi:10.1186/s13068-020-01681-4

Schindelin J, Arganda-Carreras I, Frise E, Kaynig V, Longair M, Pietzsch T, Preibisch S, Rueden C, Saalfeld S, Schmid B, Tinevez J-Y, White DJ, Hartenstein V, Eliceiri K, Tomancak P, Cardona A. 2012. Fiji: an open-source platform for biological-image analysis. Nat Methods 9:676–682. doi:10.1038/nmeth.2019

Scholz MJ, Weiss TL, Jinkerson RE, Jing J, Roth R, Goodenough U, Posewitz MC, Gerken HG. 2014. Ultrastructure and Composition of the *Nannochloropsis gaditana* Cell Wall. Eukaryot Cell 13:1450–1464. doi:10.1128/EC.00183-14

Ševčíková T, Yurchenko T, Fawley KP, Amaral R, Strnad H, Santos LMA, Fawley MW, Eliáš M. 2019. Plastid Genomes and Proteins Illuminate the Evolution of Eustigmatophyte Algae and Their Bacterial Endosymbionts. Genome Biol Evol 11:362–379. doi:10.1093/gbe/evz004

Shi J, Cui M, Yang L, Kim Y-J, Zhang D. 2015. Genetic and Biochemical Mechanisms of Pollen Wall Development. Trends Plant Sci 20:741–753. doi:10.1016/j.tplants.2015.07.010

Shimokawara M, Nishimura M, Matsuda T, Akiyama N, Kawai T. 2010. Bound forms, compositional features, major sources and diagenesis of long chain, alkyl mid-chain diols in Lake Baikal sediments over the past 28,000 years. Org Geochem 41:753–766. doi:10.1016/j.orggeochem.2010.05.013

Simionato D, Block MA, La Rocca N, Jouhet J, Maréchal E, Finazzi G, Morosinotto T. 2013. The Response of Nannochloropsis gaditana to Nitrogen Starvation Includes De Novo Biosynthesis of Triacylglycerols, a Decrease of Chloroplast Galactolipids, and Reorganization of the Photosynthetic Apparatus. Eukaryot Cell 12:665–676. doi:10.1128/ec.00363-12

Suda S, Atsumi M, Miyashita H. 2002. Taxonomic characterization of a marine *Nannochloropsis* species, *N. oceanica* sp. nov. (Eustigmatophyceae). Phycologia 41:273–279. doi:10.2216/i0031-8884-41-3-273.1

Südfeld C, Hubáček M, D’Adamo S, Wijffels RH, Barbosa MJ. 2021. Optimization of high-throughput lipid screening of the microalga *Nannochloropsis oceanica* using BODIPY 505/515. Algal Res 53:102138. doi:10.1016/j.algal.2020.102138

Sukenik A. 1990. Ecophysiological Considerations in the Optimization of Eicosapentaenoic Acid Production by Nannochloropsissp. (Eustigmatophyceae). Bioresour Technol 35:263–269.

Sukenik A, Carmeli Y. 1990. Lipid Synthesis and Fatty Acid Composition in *Nannochloropsis* sp. (Eustigmatophyceae) Grown in a Light-Dark Cycle. J Phycol 26:463–469. 10.1111/j.0022-3646.1990.00463.x

Thiam AR, Ikonen E. 2021. Lipid Droplet Nucleation. Trends Cell Biol 31:108–118. doi:10.1016/j.tcb.2020.11.006

Ueki N, Ide T, Mochiji S, Kobayashi Y, Tokutsu R, Ohnishi N, Yamaguchi K, Shigenobu S, Tanaka K, Minagawa J, Hisabori T, Hirono M, Wakabayashi K. 2016. Eyespot-dependent determination of the phototactic sign in *Chlamydomonas reinhardtii*. Proc Natl Acad Sci 113:5299–5304. doi:10.1073/pnas.1525538113

Vandenbroucke M, Largeau C. 2007. Kerogen origin, evolution and structure. Org Geochem 38:719–833. doi:10.1016/j.orggeochem.2007.01.001

Varela JC, Pereira H, Vila M, León R. 2015. Production of carotenoids by microalgae: achievements and challenges. Photosynth Res 125:423–436. doi:10.1007/s11120-015-0149-2

Veldhuis MJW, Cucci TL, Sieracki ME. 1997. Cellular DNA Content of Marine Phytoplankton Using Two New Fluorochromes: Taxonomic and Ecological Implications. J Phycol 33:527–541. 10.1111/j.0022-3646.1997.00527.x

Vermaas WFJ, Timlin JA, Jones HDT, Sinclair MB, Nieman LT, Hamad SW, Melgaard DK, Haaland DM. 2008. In vivo hyperspectral confocal fluorescence imaging to determine pigment localization and distribution in cyanobacterial cells. Proc Natl Acad Sci 105:4050–4055. doi:10.1073/pnas.0708090105

Vieler A, Wu G, Tsai C-H, Bullard B, Cornish AJ, Harvey C, Reca I-B, Thornburg C, Achawanantakun R, Buehl CJ, Campbell MS, Cavalier D, Childs KL, Clark TJ, Deshpande R, Erickson E, Ferguson AA, Handee W, Kong Q, Li X, Liu B, Lundback S, Peng C, Roston RL, Sanjaya, Simpson JP, TerBush A, Warakanont J, Zäuner S, Farre EM, Hegg EL, Jiang N, Kuo M-H, Lu Y, Niyogi KK, Ohlrogge J, Osteryoung KW, Shachar-Hill Y, Sears BB, Sun Y, Takahashi H, Yandell M, Shiu S-H, Benning C. 2012. Genome, Functional Gene Annotation, and Nuclear Transformation of the Heterokont Oleaginous Alga *Nannochloropsis oceanica* CCMP1779. PLOS Genet 8:e1003064. doi:10.1371/journal.pgen.1003064

Wang D, Ning K, Li J, Hu J, Han D, Wang H, Zeng X, Jing X, Zhou Q, Su X, Chang X, Wang A, Wang W, Jia J, Wei L, Xin Y, Qiao Y, Huang R, Chen J, Han B, Yoon K, Hill RT, Zohar Y, Chen F, Hu Q, Xu J. 2014. *Nannochloropsis* Genomes Reveal Evolution of Microalgal Oleaginous Traits. PLoS Genet 10:e1004094. doi:10.1371/journal.pgen.1004094

Wang Q, Lu Y, Xin Y, Wei L, Huang S, Xu J. 2016. Genome editing of model oleaginous microalgae *Nannochloropsis* spp. by CRISPR/Cas9. Plant J 88:1071–1081. 10.1111/tpj.13307

Wang X, Wei H, Mao X, Liu J. 2019. Proteomics Analysis of Lipid Droplets from the Oleaginous Alga *Chromochloris zofingiensis* Reveals Novel Proteins for Lipid Metabolism. Genomics Proteomics Bioinformatics 17:260–272. doi:10.1016/j.gpb.2019.01.003

Wolf FT, Stevens MV. 1967. The Fluorescence of Carotenoids. Photochem Photobiol 6:597–599. 10.1111/j.1751-1097.1967.tb08761.x

Ye Y, Huang J-C. 2020. Defining the biosynthesis of ketocarotenoids in *Chromochloris zofingiensis*. Plant Divers 2:61–66. doi:10.1016/j.pld.2019.11.001

Yurchenko T, Ševčíková T, Přibyl P, El Karkouri K, Klimeš V, Amaral R, Zbránková V, Kim E, Raoult D, Santos LMA, Eliáš M. 2018. A gene transfer event suggests a long-term partnership between eustigmatophyte algae and a novel lineage of endosymbiotic bacteria. ISME J 12:2163–2175. doi:10.1038/s41396-018-0177-y

Yurchenko T, Ševčíková T, Strnad H, Butenko A, Eliáš M. 2016. The plastid genome of some eustigmatophyte algae harbours a bacteria-derived six-gene cluster for biosynthesis of a novel secondary metabolite. Open Biol 6. doi:10.1098/rsob.160249

Zhang Z, Volkman JK. 2017. Algaenan structure in the microalga *Nannochloropsis oculata* characterized from stepwise pyrolysis. Org Geochem 104:1–7. doi:10.1016/j.orggeochem.2016.11.005

Zienkiewicz A, Zienkiewicz K, Poliner E, Pulman JA, Du Z-Y, Stefano G, Tsai C-H, Horn P, Feussner I, Farre EM, Childs KL, Brandizzi F, Benning C. 2020. The Microalga *Nannochloropsis* during Transition from Quiescence to Autotrophy in Response to Nitrogen Availability. Plant Physiol 182:819–839. doi:10.1104/pp.19.00854

Zych M, Burczyk J, Kotowska M, Kapuścik A, Banaś A, Stolarczyk A, Termińska-Pabis K, Dudek S, Klasik S. 2009. Differences in staining of the unicellular algae Chlorococcales as a function of algaenan content. Acta Agron Hung 57:377–381. doi:10.1556/AAgr.57.2009.3.12

